# Assessing the Sustainability of Pacific Walrus Harvest in a Changing Environment

**DOI:** 10.1101/2024.05.29.596516

**Authors:** Devin L. Johnson, Joseph M. Eisaguirre, Rebecca L. Taylor, Erik M. Andersen, Joel L. Garlich-Miller

## Abstract

Harvest sustainability is a primary goal of wildlife management and conservation, and in a changing world it is increasingly important to consider environmental drivers of population dynamics alongside harvest in cohesive management plans. This is particularly pertinent for harvested species that are acutely experiencing effects of climate change. The Pacific walrus (*Odobenus rosmarus divergens*), a critical traditional subsistence resource for indigenous communities, is simultaneously subject to rapid habitat loss associated with diminishing sea ice and an increasing anthropogenic footprint in the Arctic. We developed a theta-logistic population modeling-management framework to evaluate various harvest scenarios combined with four potential climate/disturbance scenarios (ranging from optimistic–pessimistic) which simulates Pacific walrus population dynamics to the end of the 21^st^ century. We considered two types of harvest strategies: (1) adaptive harvest scenarios wherein harvest is calculated as a percentage of the population and annual harvests are updated at set intervals as the population is reassessed, and (2) non-adaptive harvest scenarios wherein annual harvest remains constant. All climate/disturbance scenarios indicated declines of varying severity in Pacific walrus abundance to the end of the 21^st^ century, even in the absence of harvest. However, we found that an adaptive annual harvest of 1.23% of the independent-aged female subset of the population (e.g., 1,280 independent-aged females harvested in 2020, representing contemporary harvest levels) met our criterion for sustainability (>70% probability of maintaining population abundance above maximum net productivity level) under all climate/disturbance scenarios, accepting a medium risk tolerance level of 25%. This suggests that the present rate of Pacific walrus harvest is sustainable and will continue to be—provided the harvest adapts to match changes in population dynamics. Our simulations suggest that a sustainable non-adaptive harvest is also possible, but only at low levels if the population declines as expected. Applying a constant annual harvest of 1,280 independent-aged females (equivalent to contemporary harvest levels of 1.23) exceeded our criterion for sustainability and resulted in a >5% chance of quasi-extinction by the end of the 21^st^ century under three of the four climate/disturbance scenarios we evaluated. Our results highlight the importance of adaptive co-management strategies, and we suggest such modeling frameworks are useful for managing for harvest sustainability in a changing climate.

## INTRODUCTION

Fish and wildlife resources comprise a major component of the global economy and provide an integral source of food and economic security to subsistence communities (e.g., Inuit Circumpolar Council-Alaska, 2015). Historically, the overexploitation of wildlife populations for commercial gain has led to widespread declines across taxa and contributed to a global loss of biodiversity in the 20^th^ century (e.g., Novachek & Cleland, 2001). While territorial and government agencies were often an integral component of early exploitation, more recent agency engagements (e.g., by the United States Fish and Wildlife Service [FWS]) have focused on developing management plans to curtail these declines and have had success in establishing sustainable harvest goals for several large mammal species (e.g., Lubow & Smith 2010). In recent years, climate change and an expanding anthropogenic footprint are causing increasingly severe and unpredictable impacts on wildlife populations and ecosystems (Post et al., 2009). Thus, incorporating the effects of climate change and anthropogenic disturbance into flexible management plans can promote sustainable harvests of wildlife populations in a changing world.

Harvest sustainability is a central pillar of effective wildlife management: the goal is to maintain wildlife populations at specified levels while allowing a level of harvest that meets the needs of stakeholder groups over a set timeframe (e.g., Weinbaum et al., 2012). The specific criteria that determine whether a harvest is sustainable vary by population and are often expressed as a set of management objectives related to that population’s abundance and growth rate, along with stakeholder needs (e.g., Regehr et al., 2017). Population modeling can forecast the impacts of different harvest scenarios on population dynamics over a set timeframe and weigh outcomes against these pre-defined management objectives in a formal harvest sustainability assessment (also referred to as a “harvest risk assessment” e.g., Johnson et al., 2018). Such assessments are valuable to wildlife managers and stakeholders and can be modified for use in a formal statistical decision theoretic framework (e.g., Williams & Hooten, 2016). Additionally, climate change and anthropogenic disturbance scenarios can be incorporated into these harvest assessments (e.g., Regehr et al., 2021a), which is particularly important for harvested species undergoing rapid climate-associated habitat loss.

The Pacific walrus (*Odobenus rosmarus divergens*) is a large, ice-associated pinniped that inhabits Arctic and sub-Arctic waters and coasts of the Chukchi and Bering seas. Sea ice is an important component of their habitat—providing a platform on which to rest between foraging bouts and a safe location to breed and birth (Fay 1982, MacCracken 2017). Arctic sea ice is rapidly declining as global temperatures rise (e.g., Overland et al., 2019), resulting in unprecedented interannual variability in sea ice availability and generally longer ice-free periods across the traditional range of the Pacific walrus (Udevitz et al., 2017). Sea ice provides resting substrate near productive offshore foraging areas, which decreases the amount of time walruses spend swimming and increases the amount of time resting and foraging (Jay et al. 2017). As sea ice has declined, females and juveniles spend more time resting on land and less time resting on ice (Fischbach et al. 2022). Walruses must expend more energy when they rest on land to swim to their foraging grounds (Udevitz et al., 2017). In addition, when walruses haulout to rest in very large aggregations on land, they are exposed to increased risk of mortality from both predation and anthropogenic disturbance (Udevitz et al., 2013). Other disturbance factors associated with an expanding anthropogenic footprint in an increasingly ice-free Arctic (e.g., oil and gas exploration, shipping lanes, and commercial fishing) are also concerns of wildlife managers and Alaska Native Indigenous Knowledge (IK) holders (i.e., members of subsistence walrus-hunting communities; MacCracken et al., 2017; Metcalf & Robards, 2008). Although the size of the Pacific walrus population is relatively large (95% Bayesian credible interval (CrI) = 171,138–366,366 individuals; Beatty et al., 2022) and thought to be stationary (Taylor et al., 2018), climate change and anthropogenic disturbance are predicted to drive population declines by the middle and end of the 21^st^ century (MacCracken 2012, MacCracken et al., 2017).

Pacific walruses are a vital subsistence resource for Indigenous communities in the Bering Strait region, and were harvested for millennia at rates thought to be sustainable (MacCracken et al., 2017). In the late 19^th^ century, commercial whalers began targeting walruses, which depleted the population by the early 1900s (Fay et al., 1989) and concurrently contributed to a large-scale famine that decimated the walrus-hunting communities of Saint Lawrence Island (Metcalf & Robards, 2008). Harvest quotas implemented in the 1960s and 1970s resulted in relatively low harvest levels (4,000–6,000 individuals/year) that allowed the population to grow (Fay et al., 1989) and exceed carrying capacity by 1981 (Taylor and Udevitz 2015, Taylor et al., 2018). A spike in walrus harvest (10,000–16,000 individuals/year) in the mid-1980s concurrent with a decrease in the proportion of reproductive-aged individuals in the population contributed to another, multi-decade population decline (Fay et al., 1989, Taylor and Udevitz 2015, Taylor et al., 2018). Calf production increased from 1990 to 2015 (Taylor and Udevitz 2015, Taylor et al. 2018), and harvest levels have been relatively low and stable for the past 20 years (e.g., 4,000– 6,000 individuals/year). The multi-decade population decline ended by the mid-2010s, when the walrus population size stabilized (Taylor et al. 2018), although the current carrying capacity may be lower than the carrying capacity that was exceeded at the outset of the last population decline (Taylor et al., 2018).

The Pacific walrus harvest has been limited to traditional subsistence users in Alaska and Chukotka since an end to the Russian commercial harvest in 1992 (Garlich-Miller et al. 2006). The Alaska harvest has been self-regulated by local Indigenous walrus-hunting communities since 1979 due to lack of regulatory authority by federal agencies to regulate harvest of a legally non-depleted population (Robards and Joly, 2007; Gadamus & Raymond-Yakoubian, 2015; MacCracken et al., 2017); whereas Chukotkan communities adhere to annual quotas set by the Russian government (Kryukova, 2019). A comanagement agreement exists between the FWS and the Eskimo Walrus Commission (EWC; a commission represented by 19 Yup’ik, St, Lawrence Island Yupik, and Iñupiaq communities) with the shared goal of conserving the Pacific walrus population while promoting a sustainable subsistence harvest (Metcalf & Robards, 2008). Specifically, one role of the FWS in this comanagement agreement is to “provide the EWC with information on walrus population, status, and trends for the development of sound management practices integral to fulfilling their mission of representing the interests of subsistence users and walrus hunters” (Metcalf & Robards, 2008). Although previous efforts by western scientists suggest that current harvest levels are sustainable (MacCracken et al., 2017; USFWS 2023), there is concern over the cumulative impacts of climate change, anthropogenic disturbance, and harvest on future Pacific walrus population dynamics.

We developed a predictive population model to evaluate the impacts of different harvest scenarios on the Pacific walrus population to the end of the 21^st^ century. Harvest scenarios were analyzed in concert with four combined scenarios that incorporated varying degrees of sea ice loss from climate change along with different anthropogenic disturbance scenarios (Table 1; Johnson et al., 2023). Our underlying model is based on a discrete version of the theta-logistic density-dependent population growth equation, which has been successfully implemented in harvest assessments for other arctic marine mammal populations (e.g., polar bears [*Ursus maritimus*]; Regehr et al., 2021). This approach is recommended when incomplete information is available on the age-sex composition of the harvest or population (Johnson et al., 2018). We focus on the independent-aged female segment of the population because females drive population dynamics in polygynous species such as walruses, more so than males. We developed harvest scenarios based on historical harvest rates, demographic trends in the harvest (Garlich-Miller et al., 2006), and IK holder input. Climate change and disturbance scenarios were developed and incorporated in the theta-logistic model parameters for carrying capacity (*K*) and the maximum intrinsic rate of population growth (*r_max_*) – both of which are expected to decline at different rates based on future sea ice conditions (MacCracken 2012, Johnson et al., 2023). We introduce a flexible, intuitive framework that can be used by managers and the subsistence hunting community to assess the sustainability and risk of different harvest scenarios for the Pacific walrus in a rapidly changing environment.

**Table 1.** Summary of climate/disturbance scenarios used to derive estimates of *K_t_* and *r_max,t_* for the independent-aged female Pacific walrus population (from the Johnson et al., 2023 PCoD [Population Consequences of Disturbance] model). In the scenarios described below, we incorporated sea ice data from two shared socioeconomic pathways (SSPs) used in global climate models: ssp245 represents an intermediate (“middle of the road”) scenario, whereas ssp585 represents a pessimistic (fossil-fueled development, “business as usual”) scenario (Fox-Kemper et al. 2021). Scenarios also incorporate varying intensities of anthropogenic disturbance (e.g., future oil & gas development, potential changes to resource quality, and terrestrial haulout mortality factors) that were considered in the PCoD model (Johnson et al., 2023). We display mid-century (2050) and end-century (2100) estimates as 95% credibility intervals for *K_t_* and *r_max,t_*.

| Name | Climate Scenario | Disturbance Intensity | $K_{2050}$ | $K_{2100}$ | $r_{max,2050}$ | $r_{max,2100}$ |
| --- | --- | --- | --- | --- | --- | --- |
| Optimistic | ssp245 | Low | 79,531–146,523 | 74,148–139,056 | 0.033–0.042 | 0.030–0.041 |
| Intermediate_245 | ssp245 | Medium | 58,993–100,178 | 58,871–90,113 | 0.031–0.040 | 0.026–0.039 |
| Intermediate_585 | ssp585 | Medium | 58,793–97,754 | 47,030–73,693 | 0.030–0.039 | 0.025–0.037 |
| Pessimistic | ssp585 | High | 39,171–63,124 | 31,959–49,805 | 0.025–0.038 | 0.020–0.033 |

## METHODS

### Model Structure & Overview

This harvest assessment uses a modified discrete-time version of the theta-logistic population growth model (following Regehr et al., 2021a), which can be expressed as the following equation:

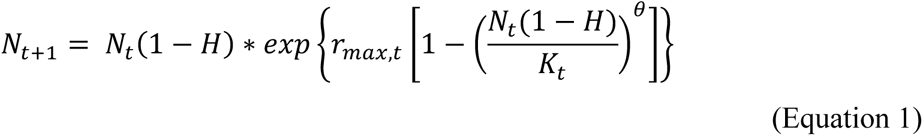

In Equation 1: *N_t_* is the population size (independent-aged females) in year *t*; *H* is the proportion of the population harvested annually; *r_max,t_* is the intrinsic growth rate of the population in year *t*; *K_t_* is the carrying capacity in year *t*; and *θ* is a shape parameter that determines how the growth rate changes as a function of density. The model focuses on the “independent-aged” female portion of the Pacific walrus population, which we define as females of post-weaning, reproductive, and post-reproductive age (>= 2 years). We define *r_max_* as the maximum intrinsic rate of population growth (e.g., Cortés 2016), which expresses how quickly the population could grow under ideal circumstances. We held *θ* constant at 5.045 because this value produces dynamics typical of long-lived mammals (Wade, 1998) and has recently been used in a harvest assessment for polar bears (Regehr et al., 2021).

This modelling effort required four major components: 1) an initial abundance estimate for the independent-aged female portion of the Pacific walrus population (*N_t = 1_*); 2) a suite of harvest scenarios based on historic subsistence use; 3) estimates of *K_t_* and *r_max,t_* for the population at each timestep of the simulation under different climate/disturbance scenarios; and 4) a set of management objectives against which to compare model output. Ultimately, we produced a total of 200 simulations to assess harvest sustainability and quasi-extinction probability under an array of harvest/climate change scenarios (Fig. 1).

**Figure 1.**
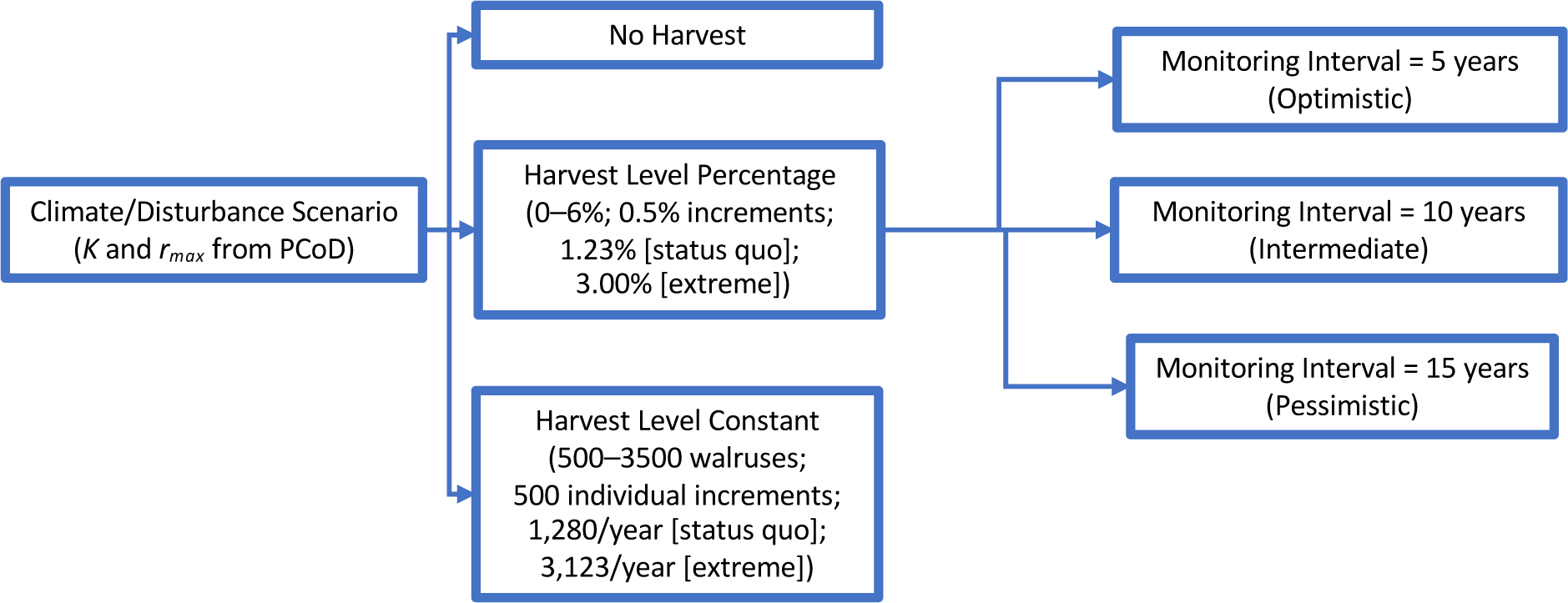
Management scenarios considered in this analysis. Four climate-disturbance scenarios (see Table 1) were implemented using the Pacific walrus population consequences of disturbance/climate change (PCoD model, Johnson et al., 2023) to produce time-specific estimates of carrying capacity (*K^t^*) and intrinsic rate of increase (*r_max,t_*), which we then used in the theta-logistic model in the present study.

### Abundance and Historical Demographic Harvest Estimates

The theta-logistic model requires an initial abundance estimate for the “independent-aged” female portion of the Pacific walrus population. Although past efforts to estimate walrus population abundance have been infrequent and resulted in large confidence intervals (Table S1), recent advances in genetic mark recapture approaches allowed for a more precise estimate that can be applied to a demographic subset of the population. Thus, we used the most recent estimate of Pacific walrus population abundance (Beatty et al., 2022) to derive an estimate of independent-aged female abundance (*N_2020_*; mean = 104,123; 95% CrI = 65,827–142,418), which was used for forward simulations in the theta-logistic model.

To construct harvest scenarios for the model, we analyzed historical harvest trends in conjunction with demographic population estimates, both of which are described below. Pacific walruses are harvested annually by traditional subsistence users in both Alaska and Russia. FWS monitoring programs indicate a high level of hunter reporting compliance (76%, Garlich-Miller & Burn, 2009) in the Alaskan communities of Gambell and Savoonga, which together account for >80% of the US walrus harvest since 1990 (MacCracken et al., 2017). Russian hunter compliance is estimated to be ∼80% (Smirnov et al., 2002). Annual harvest estimates were compiled and adjusted to account for a 42% struck-and-lost rate (Fay et al., 1994) and underreporting as described above. Total estimated harvest in recent years (2005–2015) has been 28% lower than mean harvest levels from 1975–2005 (Fig. 2A), and it is hypothesized that changes in sea ice dynamics may be limiting subsistence hunter access and success rates (Hovelsrud, 2008). Since the 1970s, the proportion of independent-aged females in the US harvest has been fairly stable (Fig. S1; Garlich-Miller et al., 2006) at approximately 31% of the overall harvest (Fig. 2). Russian communities harvested a lower proportion of independent-aged females (approximately 22% of their overall harvest; Fig. 2), and a limited demographic dataset indicates that there has been no significant temporal trend in the Russian harvest (Fig. S1; Kryukova, 2019).

**Figure 2.**
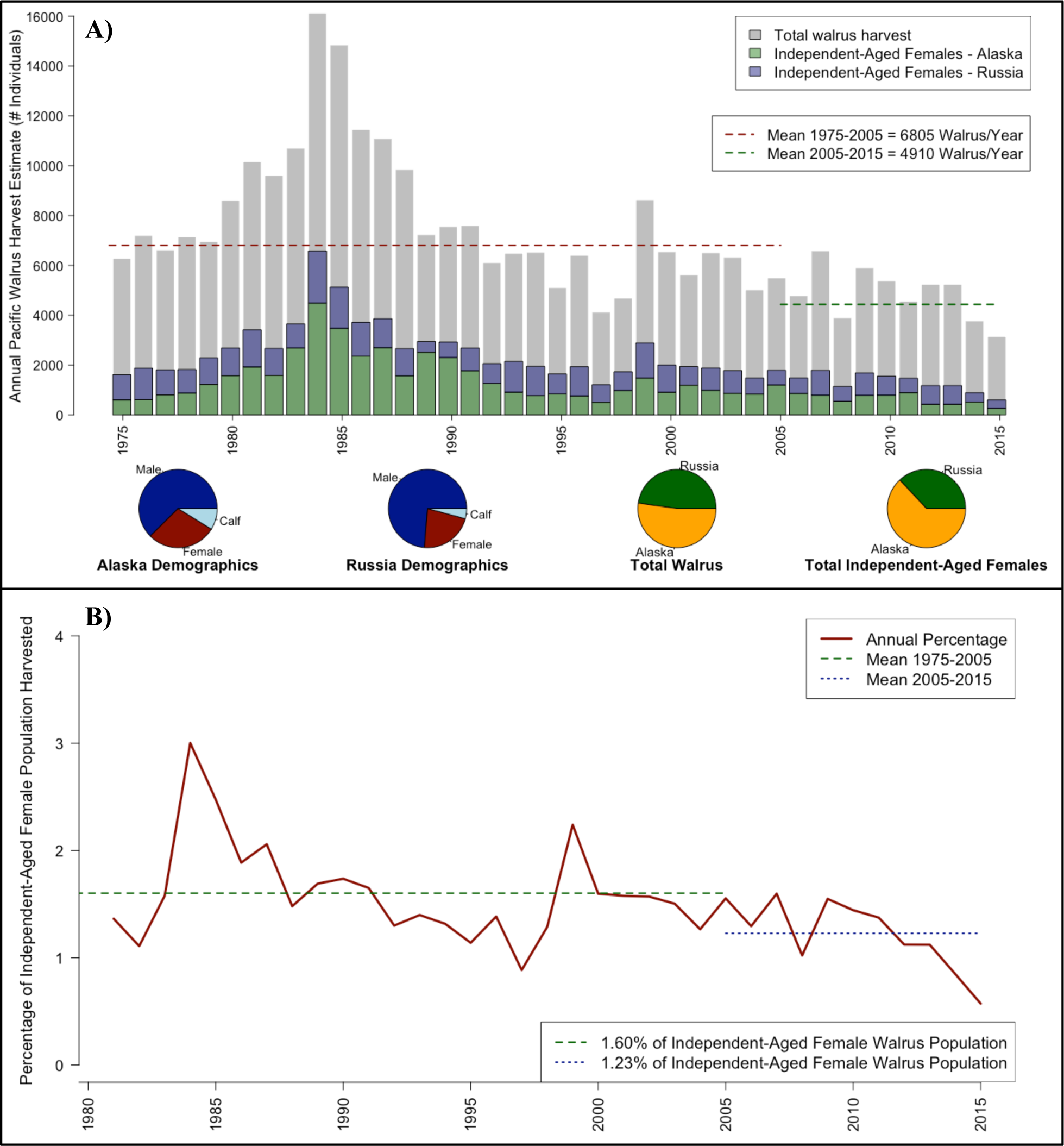
A) Pacific walrus harvest demographics 1975-2015; and B) the estimated percentage of the independent-aged female population that was harvested in each year based on the harvest data found in Fig. 1A scaled to absolute population size (Beatty et al. 2022) and population trend and age structure estimates (Taylor et al., 2018). In panel A, gray bars indicate total overall mean harvest numbers (Alaska and Russia combined; adjusted for hunter reporting compliance and struck-and-loss rates), and colored bars indicate the estimated numbers of independent-aged females harvested in each year. Pie charts indicate the estimated demographic breakdown from 2005-2015 for Alaska and Russia, and the regional breakdown of total and independent-aged female harvest. In panel B, the green dashed line indicates the mean harvest rate from 1975-2005, and the blue dotted line indicates the mean harvest rate from 2005-2015.

To determine the proportion of the independent-aged female population that has been historically harvested, we estimated independent-aged female abundance from 1975–2015 (Fig. 2B). We combined the estimated population trend in the independent-aged female segment of the population from 1975–2015 (Taylor et al., 2018) with an estimate of the absolute size of the independent-aged female population in 2013–2017 (Beatty et al., 2022). Taylor et al. (2018) reported a significant decline from 1975–2015 both in Pacific walrus population size and in the proportion of the independent-aged female population; our analyses incorporated these trends. We determined the proportion of the independent-aged females harvested in each year of the historical datasets for both Alaska and Russia based on that year’s estimates of number of independent-aged females harvested and number of independent-aged females in the population. This process assumes that demographic harvest data for the recorded communities is representative of overall demographic harvest trends for the country they represent. The resulting annual harvest percentages (Fig. 2B) represent historical and more contemporary estimates of *H* (i.e., the percentage of the independent-aged female population that was harvested in each year) which were used to inform harvest scenarios.

### Incorporating Indigenous Knowledge

This model was developed to inform a co-management framework, and IK holders were included in various stages of the model’s development. Specifically, a steering committee consisting of agency biologists (FWS & USGS) and members of the EWC were consulted on the general methodological approach and developed the harvest scenarios and management objectives addressed in this study. Additionally, the steering committee identified the need for further consultation and discussion with members of walrus-hunting communities to gather IK on walrus behavior and harvest patterns in the face of climate change. In August 2023, a 3-day IK workshop (Metcalf et al. 2023) was held where agency biologists (DLJ & JLGM) discussed the harvest assessment model with five experienced walrus hunters from the villages of Gambell and Savoonga to gather additional information related to the future of harvest, and ultimately provide input on relevant harvest scenarios and management objectives to be assessed using the model framework.

### Harvest Scenarios

We implemented a theta-logistic model designed to assess the response of independent-aged female population size to different harvest rates, (*H*, the proportion of the independent-aged female Pacific walrus population that is harvested annually). We used values of *H* ranging from 0–6% by increments of 0.5%. Along with these generalized values, we built scenarios with more specific *H* values based on the historical harvest. We considered an annual harvest rate of 1.23% of the independent-aged female population to represent the “status quo” harvest level for contemporary conditions (Fig. 2B), because it represents the mean harvest level from 2005– 2015. We considered an annual harvest rate of 3.00% to represent an “extreme harvest” scenario, which represents the highest annual harvest rate in the recent historical record (i.e., 1984; Fig. 2).

Because *H* is expressed as a proportion of the population, the number of walruses harvested in adaptive scenarios changes in response to projected changes in population size. Pacific walrus abundance has historically been estimated infrequently (Table S1) due to logistical constraints. Thus, the simulations include a “monitoring interval” term that expresses how frequently abundance is estimated and used to update the harvest quota into the future. We considered 5-year, 10-year, and 15-year monitoring intervals in our simulations, which represent the range of historical population reassessment intervals (Table S1) and are in line with FWS planned population assessments.

Finally, using this framework, we can simulate an annual harvest that is held constant regardless of projected changes to population size. This may represent a situation where management agencies are unable to adequately reassess population abundance, or a scenario where walrus-hunting communities have a set minimum number of individuals they need to harvest annually to meet community needs. These scenarios use the theta-logistic equation with a slight modification as follows:

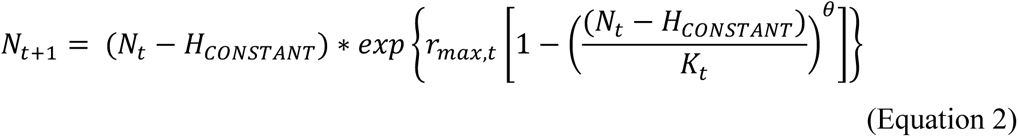

We included scenarios with constant number of harvested animals ranging from 500–2,500 independent-aged females (by 500-individual increments), which approximate the range of independent-aged females harvested historically (Fig. 2). Thus, the scenarios with a constant number of harvested animals represent scenarios with a changing harvest rate because while the number of harvest animals remains constant, population abundance will change over time.

### Climate Change and Disturbance Scenarios

Pacific walrus carrying capacity (*K_t_*) and maximum intrinsic population growth (*r_max,t_*) form the basis for population growth or decline in the theta-logistic model, and they are expected to change in relation to declining sea ice availability and other anthropogenic factors. The degree of change can be predicted using the population consequences of disturbance/climate change (PCoD) framework developed by Johnson et al. (2023). Briefly, the PCoD framework implements a state-dependent bioenergetic model that simulates independent-aged female body condition and reproductive state, linking exposure to various stressors (e.g., reduced access to sea ice, disturbance from oil development) to changes in body condition and ultimately population-level parameters (i.e., *K_t_* and *r_max,t_*). The PCoD model was designed to incorporate sea ice projections to the middle and end of the 21^st^ century under two different global climate scenarios (ssp245 [middle of the road]; ssp585 [fossil-fueled development]; Fox-Kemper et al., 2021), alongside various anthropogenic disturbance scenarios. Johnson et al. (2023) used input from wildlife managers and IK holders to develop a set of four combined climate change/disturbance scenarios that are particularly relevant to harvest management. They then used the PCoD model to produce estimates of *K_t_* and *r_max,t_* in 2050 and 2100 under those scenarios, and we use these estimates in the present study (Table 1; Fig. S2).

### Simulations

We used the theta-logistic model (Equations 1 & 2) to simulate Pacific walrus population dynamics under a suite of climate/disturbance scenarios (Table 1) and harvest scenarios (Fig. 1). For each climate/disturbance scenario, we simulated a total of 50 different harvest scenarios. Each simulation was comprised of 1,000 population projections, each of which was based on a random draw from the full posterior distributions of input parameters. We expressed simulation output as 95% credibility intervals (i.e., the interval within which 95% of 1,000 projections fell for each simulation).

Independent-aged female walrus abundance in 2020 (N_2020_) was drawn from the posterior distribution of the independent-aged female population size estimate derived from Beatty et al., 2022 (mean = 104,123; 95% CrI = 65,827–142,418), which applies to the years 2013–2017. For ease of interpretation, our simulations run from the years 2020–2100; thus, we assumed population abundance and harvest patterns did not markedly change between the years 2017– 2020. We felt our assumption was appropriate because walruses have high juvenile and adult survival coupled with a low reproductive rate. Distributions for *K_t_* and *r_max,t_* were generated using the Pacific Walrus PCoD framework (Johnson et al., 2023) and are specific to the four climate/disturbance scenarios. Because the PCoD model produces distributions of *K_t_* and *r_max,t_* for the middle and end of the 21^st^ century, we developed a sampling protocol to estimate each of these parameters in each year of the harvest sustainability simulation. Distributions for each parameter were generated from 100 simulated populations (each of 100 individuals) within the PCoD framework for each climate-disturbance scenario (Fig. S2). For each parameter, we sampled the initial distribution (*K*_2020_ & *r_max,_*_2020_), calculated the slope to a sample from the intermediate distribution (*K*_2050_ & *r_max,_*_2050_) and repeated this process from the intermediate to end distribution (*K*_2100_ & *r_max,_*_2100_). We repeated this process for each population simulated with the theta-logistic model as a way of estimating *K_t_* and *r_max,t_* in each year while best accounting for uncertainty.

### Harvest Management Objectives

We worked closely with wildlife managers and IK holders to develop two management objectives for the Pacific walrus population. Management Objective 1 (MO1) was to maintain the population size above the Maximum Net Productivity Level (MNPL), which occurs approximately when *N_t_/K_t_* = 0.70 if *θ* = 5.045 for marine mammal stocks (Ragen, 1995; Regehr et al., 2021). This objective would theoretically allow for the population’s maximum sustainable yield to be harvested while avoiding overexploitation (Wade, 1998). Management Objective 2 (MO2) was to maintain the population above a pre-defined quasi-extinction threshold. For illustration, we define the quasi-extinction threshold as 5,000 independent-aged females.

Although little information exists to indicate when the population might experience permanent deleterious small-population effects (e.g., inbreeding depression), this estimate is within the range of proposed estimates of minimum viable population size for similarly long-lived mammal species (e.g., Reed et al., 2003). The probability of achieving each management objective was assessed for each simulation. Because of the wide credible interval for the initial population size, some simulated *N_2020_/K_2020_* values were < 0.70; therefore, we assessed management objectives beginning on year 2030 to allow for an appropriate equilibration interval.

Each management objective was assessed according to a pre-defined risk tolerance level. The selection of a risk tolerance level is an important component of evaluating a harvest strategy and should involve a coordinated decision-making process between managers and stakeholders. For demonstration purposes, we selected a “medium” risk tolerance level of 25% (i.e., accepting a 25% probability of failing to meet the management objective) for MO1 (sustainability), which is consistent with similar analyses on marine mammals (e.g., Regehr et al., 2021a; Regehr et al., 2021b). To characterize the sensitivity of the model to the choice of a risk tolerance level, we compared model output at a 15% (low) and 35% (high) risk tolerance level as well for several simulations (see Results: Risk Tolerance Sensitivity). We defined a comparatively low risk tolerance level of 5% for MO2 (quasi-extinction), as it represents a more severe outcome than an unsustainable harvest.

### Potential Biological Removal

In the United States, sustainable harvest of marine mammal stocks (including the Pacific walrus), is often assessed by calculating the Potential Biological Removal (PBR; Wade 1998; NMFS 2016). There is question whether the PBR equation is appropriate for species undergoing rapid habitat change (e.g., Robards et al., 2009), and whether a more sophisticated modeling approach should be used for sustainability assessments in the future. We calculated PBR for the independent-aged female subset of the Pacific walrus population to serve as a generalized metric of sustainability against which to compare model output from the sustainability analysis. PBR was calculated using parameter values from the most recent Pacific walrus stock assessment report (USFWS, 2023) and a population abundance estimate based on Beatty et al., 2022 (Supplemental Material 1).

## RESULTS

### Summary

We simulated 50 harvest scenarios for each of four climate/disturbance scenarios (characterized by specific values of *K_t_* and *r_max,t_*) for a total of 200 simulations. Each harvest scenario was characterized by each combination of two input parameters: harvest (as an adaptive rate of 0–6% by 0.5% increments (and also status quo [1.23%] harvest rates) and as a non-adaptive level of 500–2,500 walruses by 500-individual increments) and monitoring interval (5, 10, and 15 years; Fig. 1; Tables S2-S5). All models consider a risk tolerance level of 25% for MO1 (sustainability) and 5% for MO2 (quasi-extinction) unless otherwise noted. Overall, model output indicated that an adaptive status quo harvest rate (representing contemporary levels of harvest) is within a sustainable range and will continue to be to the end of the 21^st^ century under the four climate/disturbance scenarios considered (Fig. 3, Table 2). An adaptive harvest rate of up to 2.25% of the independent-aged female Pacific walrus population met the criteria for both MO1 (sustainability) and MO2 (quasi-extinction; Table S6), whereas an adaptive harvest rate of up to 5.5–6% met the criteria for MO2 but not MO1 (Fig. 3). Non-adaptive harvest simulation output indicated that a sustainable non-adaptive harvest is possible, but only at relatively low levels of harvest (i.e., under 1500 independent-aged females/year; Fig. 4). We rely on a set of examples taken from the broader set of simulations to conceptualize and discuss portions of the results in further detail.

**Figure 3.**
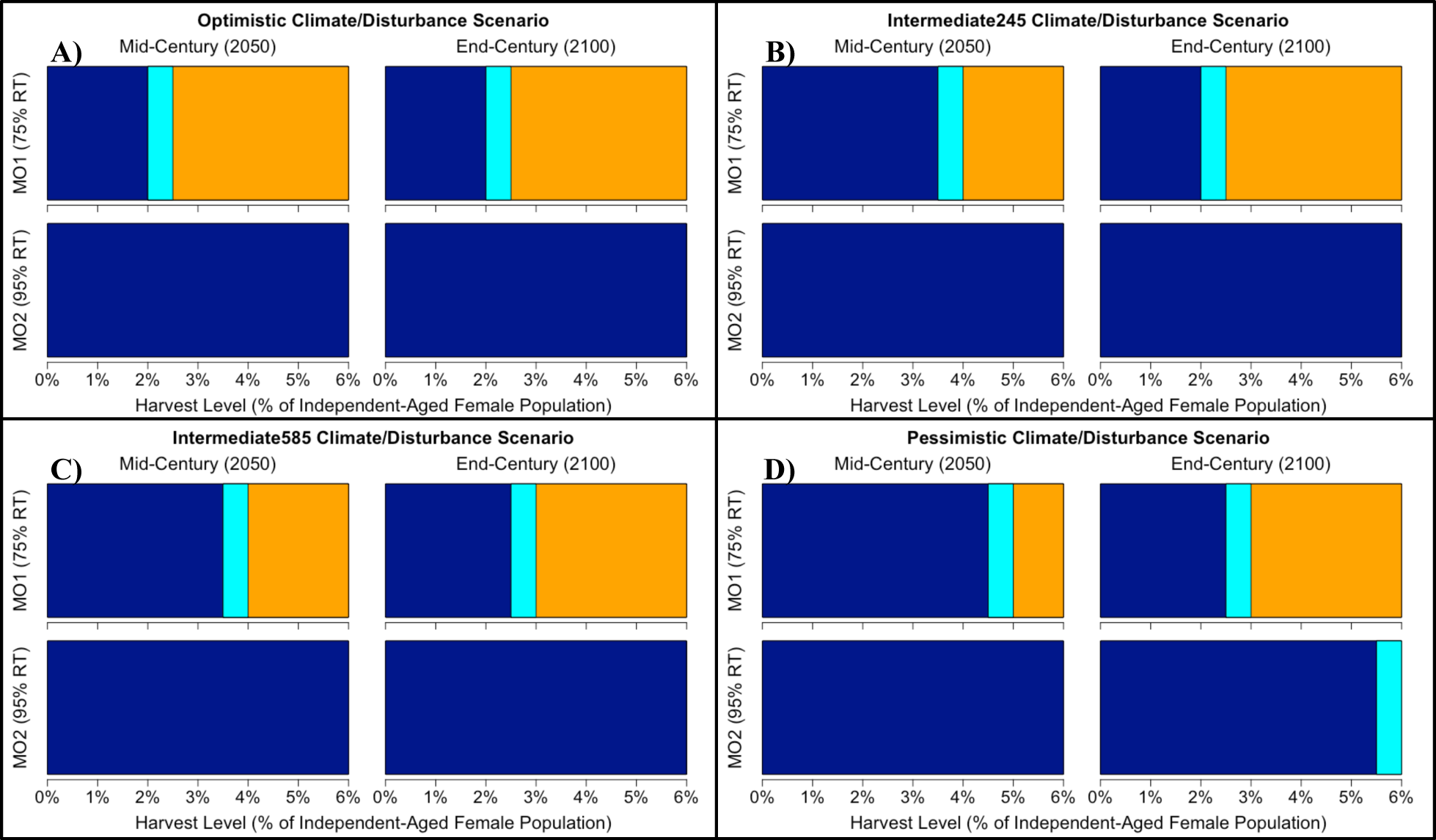
Summary of adaptive harvest scenarios, indicating the percentage-based levels of harvest that meet (dark blue) or fail to meet (orange) the two management objectives in the years 2050 and 2100 under each of four climate/disturbance scenarios (A-D). Cyan bars indicate the threshold range (in 0.5% increments) where each management objective fails. All displayed model output considers a 10-year (intermediate) monitoring interval. Management Objective 1 (MO1; harvest sustainability) was met if 750 of the 1000 projections in each scenario (75%) resulted in a population at or above the Maximum Net Productivity Level (MNPL). Management Objective 2 (MO2; quasi-extinction) was met if the independent-aged female population size was > 5000 in 950 of the 1000 projections (95%). In some cases, the non-optimistic scenarios allow for a greater mid-century sustainable harvest than the optimistic scenarios. This results from a faster decline in carrying capacity than population size in the non-optimistic scenario, which results in a surplus of animals that could be harvested (i.e., N/K > 1; e.g., Fig. 6).

**Figure 4.**
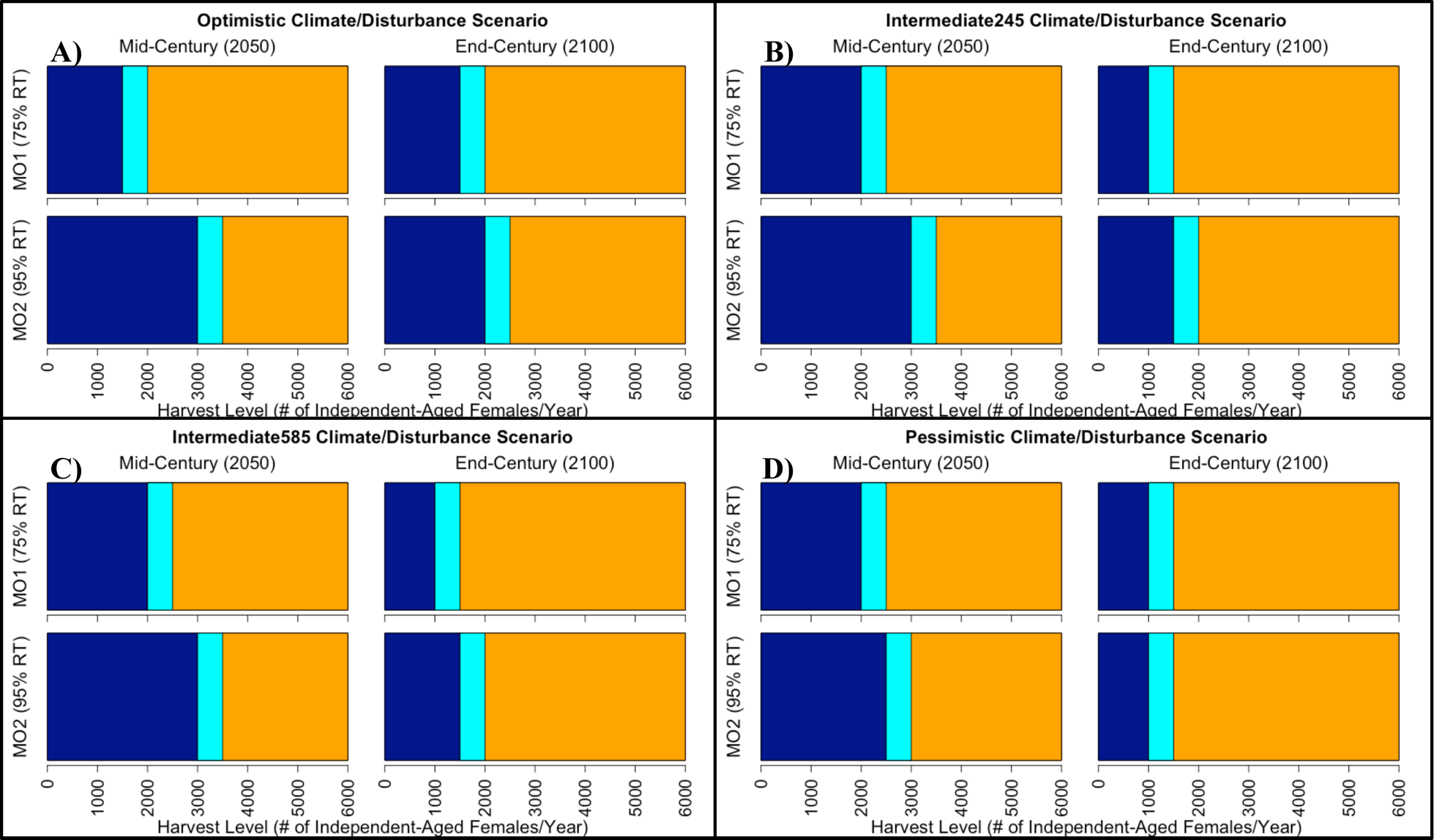
Summary of non-adaptive harvest scenarios for Pacific walrus, indicating the constant annual levels of harvest that meet (dark blue) or fail to meet (orange) the two management objectives in the years 2050 and 2100 under each of four climate/disturbance scenarios (A-D). Cyan bars indicate the threshold range (in 500-individual increments) where each management objective fails. All displayed model output considers a 10-year (intermediate) monitoring interval. Management Objective 1 (MO1; harvest sustainability) was met if 750 of the 1000 projections in each scenario (75%) resulted in a population at or above the Maximum Net Productivity Level (MNPL). Management Objective 2 (MO2; quasi-extinction) was met if the independent-aged female population size was > 5000 in 950 of the 1000 projections (95%). In some cases, the non-optimistic scenarios allow for a greater mid-century sustainable harvest than the optimistic scenarios. This results from a faster decline in carrying capacity than population size in the non-optimistic scenario, which results in a surplus of animals that could be harvested (i.e., N/K > 1; e.g., Fig. 6).

**Table 2.** Summary of model output for example scenarios seen in the Results section. Scenario is one of four climate/disturbance scenarios for *K* and *r_max_* (OPT=Optimistic, I_245=Intermediate_245, I_585=Intermediate_585, PESS=Pessimistic); H_type (harvest type) is either A (adaptive scenarios) or N-A (non-adaptive scenarios); M_int is the monitoring interval; H is the percentage of the independent-aged female Pacific walrus population that is harvested annually; A_t_ is the absolute harvest under each scenario; N_t_ is the mean estimated population size at each timestep of the simulation (i.e., the years 2050 and 2100); MO1 and MO2 are the probabilities of meeting Management Objective 1 (sustainability) and Management Objective 2 (avoiding quasi-extinction).

| Model Parameters |  |  |  |  | Output – Timestep 30 (2050) |  |  |  | Output – Timestep 80 (2100) |  |  |  |
| --- | --- | --- | --- | --- | --- | --- | --- | --- | --- | --- | --- | --- |
| Scenario | H_type | M_int | H | $A_{2020}$ | $N_{2050}$ | $A_{2050}$ | MO1<sub>2050</sub> | MO2<sub>2050</sub> | $N_{2100}$ | $A_{2100}$ | MO1<sub>2100</sub> | MO2<sub>2100</sub> |
| **Examples 1 & 2 for Adaptive Harvest: Effect of H and Effect of Climate/Disturbance Scenario** |  |  |  |  |  |  |  |  |  |  |  |  |
| OPT | A | 10 | 0.00% | 0 | 113,734 | 0 | 0.108 | 0.000 | 107,559 | 0 | 0.114 | 0.000 |
| OPT | A | 10 | 1.23% | 1,280 | 106,916 | 1,315 | 0.129 | 0.000 | 99,976 | 1,229 | 0.147 | 0.000 |
| OPT | A | 10 | 3.00% | 3,123 | 87,695 | 2,630 | 0.341 | 0.000 | 79,414 | 2,382 | 0.423 | 0.000 |
| I_245 | A | 10 | 0.00% | 0 | 84,027 | 0 | 0.054 | 0.000 | 74,234 | 0 | 0.083 | 0.000 |
| I_245 | A | 10 | 1.23% | 1,280 | 85,239 | 1,048 | 0.070 | 0.000 | 69,135 | 850 | 0.119 | 0.000 |
| I_245 | A | 10 | 3.00% | 3,123 | 69,726 | 2,227 | 0.197 | 0.000 | 53,255 | 1,597 | 0.479 | 0.000 |
| I_585 | A | 10 | 0.00% | 0 | 83,471 | 0 | 0.054 | 0.000 | 63,685 | 0 | 0.060 | 0.000 |
| I_585 | A | 10 | 1.23% | 1,280 | 84,043 | 1,033 | 0.082 | 0.000 | 59,103 | 726 | 0.107 | 0.000 |
| I_585 | A | 10 | 3.00% | 3,123 | 68,651 | 2,201 | 0.188 | 0.000 | 46,585 | 1,397 | 0.383 | 0.000 |
| PESS | A | 10 | 0.00% | 0 | 59,422 | 0 | 0.031 | 0.000 | 42,812 | 0 | 0.067 | 0.000 |
| PESS | A | 10 | 1.23% | 1,280 | 66,615 | 819 | 0.037 | 0.000 | 38,899 | 478 | 0.101 | 0.000 |
| PESS | A | 10 | 3.00% | 3,123 | 52,813 | 1,584 | 0.068 | 0.000 | 29,553 | 886 | 0.510 | 0.000 |
| **Example 3: Non-Adaptive vs Adaptive Harvest Regimes** |  |  |  |  |  |  |  |  |  |  |  |  |
| I_245 | N-A | NA | - | 1,280 | 83,847 | 1,280 | 0.109 | 0.000 | 62,831 | 1,280 | 0.288 | 0.002 |
| I_245 | N-A | NA | - | 3,123 | 60,225 | 3,123 | 0.452 | 0.041 | 3,220 | 3,123 | 0.977 | 0.915 |
| PESS | N-A | NA | - | 1,280 | 65,820 | 1,280 | 0.053 | 0.000 | 17,561 | 1,280 | 0.746 | 0.299 |
| PESS | N-A | NA | - | 3,123 | 45,853 | 3,123 | 0.397 | 0.073 | 0 | 3,123 | 1.000 | 1.000 |
| **Example 4: Monitoring Interval Sensitivity** |  |  |  |  |  |  |  |  |  |  |  |  |
| I_245 | A | 5 | 3.00% | 3,123 | 70,890 | 2,126 | 0.176 | 0.000 | 54,051 | 1,621 | 0.448 | 0.000 |
| I_245 | A | 10 | 3.00% | 3,123 | 69,726 | 2,227 | 0.197 | 0.000 | 53,255 | 1,597 | 0.479 | 0.000 |
| I_245 | A | 15 | 3.00% | 3,123 | 68,347 | 2,178 | 0.233 | 0.000 | 52,307 | 1,597 | 0.521 | 0.000 |
| I_245 | A | 5 | 4.00% | 4,164 | 61,059 | 2,442 | 0.375 | 0.000 | 38,086 | 1,523 | 0.906 | 0.000 |
| I_245 | A | 10 | 4.00% | 4,164 | 58,975 | 2,538 | 0.431 | 0.000 | 35,483 | 1,419 | 0.945 | 0.000 |
| I_245 | A | 15 | 4.00% | 4,164 | 56,816 | 2,449 | 0.473 | 0.000 | 32,979 | 1,393 | 0.967 | 0.000 |
| I_245 | A | 5 | 5.00% | 5,206 | 48,439 | 2,421 | 0.704 | 0.000 | 20,161 | 1,008 | 1.000 | 0.000 |
| I_245 | A | 10 | 5.00% | 5,206 | 43,679 | 2,438 | 0.836 | 0.000 | 15,830 | 791 | 1.000 | 0.000 |
| I_245 | A | 15 | 5.00% | 5,206 | 37,851 | 2,156 | 0.939 | 0.000 | 11,199 | 644 | 1.000 | 0.012 |
| I_245 | A | 5 | 6.00% | 6,247 | 34,195 | 2,051 | 0.977 | 0.000 | 8,274 | 496 | 1.000 | 0.022 |
| I_245 | A | 10 | 6.00% | 6,247 | 27,446 | 1,978 | 0.999 | 0.000 | 4,671 | 280 | 1.000 | 0.572 |
| I_245 | A | 15 | 6.00% | 6,247 | 18,221 | 1,424 | 1.000 | 0.000 | 1,486 | 120 | 1.000 | 0.999 |

### Effect of *H* in an Adaptive Harvest

To demonstrate the simulated effects of adaptive harvest on Pacific walrus population dynamics in isolation, we focus on three example adaptive harvest scenarios (considering an intermediate 10-year monitoring interval and using *K_t_* and *r_max,t_* from the Intermediate_245 climate/disturbance scenario): a no-harvest scenario (0%); a status-quo scenario (1.23%) that represents a contemporary harvest rate; and an “extreme harvest” scenario (3.00%) which represents the highest rate of harvest the independent-aged female portion of the population underwent on historical record (in the mid-1980s; Fig. 2B). Figure 5 illustrates the model output of these three harvest scenarios. While an adaptive status-quo harvest met our criteria for sustainability (Fig. 5B) and generally had little effect on population abundance, the extreme harvest scenario contributed to a significant population decline by the end of the 21^st^ century (*N_2100_* = 53,255 independent-aged females [51% of the contemporary abundance estimate]) and failed MO1 (sustainability) 43 years into the simulation (Fig. 5C, Table 2).

**Figure 5.**
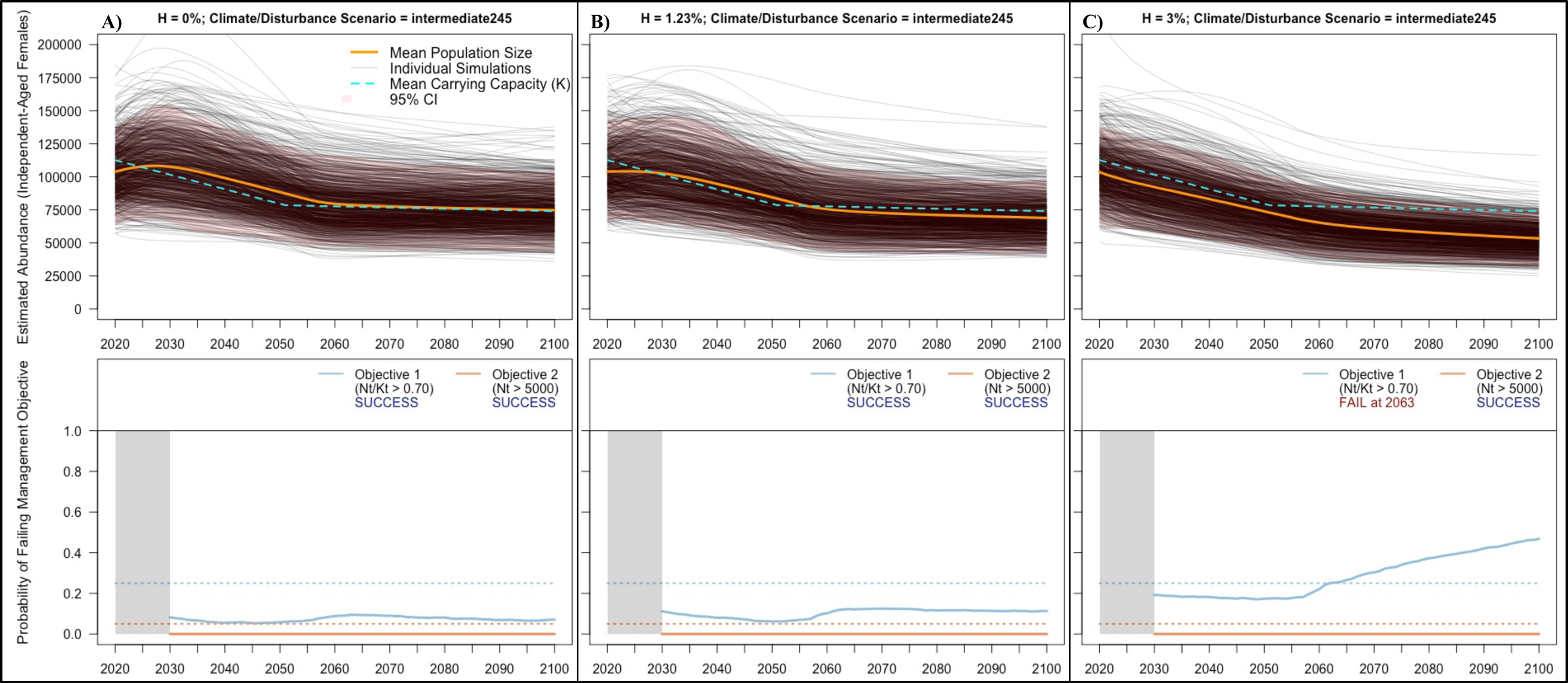
Example output from the theta-logistic model, considering three adaptive harvest scenarios under the intermediate_245 combined climate/disturbance scenario and a 10-year (intermediate) monitoring interval projecting forward to the end of the 21^st^ century. Column A considers no harvest, column B considers an annual harvest rate of 1.72% of the independent-aged female population (the mean harvest from 2005-2015), and column C considers an annual harvest rate of 3.10% of the independent-aged female population (i.e., the maximum estimated percentage-based harvest since 1975). In the top panels, orange lines represent the mean estimated population size of independent-aged female Pacific walruses, gray lines are individual simulations (n=1000), cyan dashed lines represent mean carrying capacity estimates from each climate/disturbance scenario, and the red shaded region is a 95% credible interval of theta-logistic simulations. Bottom panels represent the associated probability of failing to meet risk tolerance thresholds for pre-defined management objectives (MO1 & MO2) at each timestep of the simulation.

### Effect of Climate/Disturbance Scenario on Adaptive Harvest

To exemplify the effect of climate change and anthropogenic disturbance on harvest sustainability, we present each of the three above example adaptive harvest scenarios under each of the four climate/disturbance scenarios (Fig. 6, Table 2). Under the most pessimistic climate/disturbance scenario the simulated population declined substantially even under a no-harvest scenario, to a mean of 42,812 independent-aged females by the end of the 21^st^ century (41% of N_2020_; Fig. 6; Table 2). Despite this decline, harvesting at the status-quo rate (1.23%) only had a moderate effect on abundance (N_2100_ = 38,899 individuals; 37% of N_2020_) and met both management objectives. An extreme harvest rate (3.00%) exacerbated the population decline (N_2100_ = 29,553; 28% of N_2020_; Fig. 6, Table 2), and was not sustainable. An adaptive harvest of 5.5% (higher than any historical harvest on record) was required to surpass the quasi-extinction threshold under a pessimistic climate/disturbance scenario (MO2; risk tolerance = 5%) by the end of the century (Table S5).

**Figure 6.**
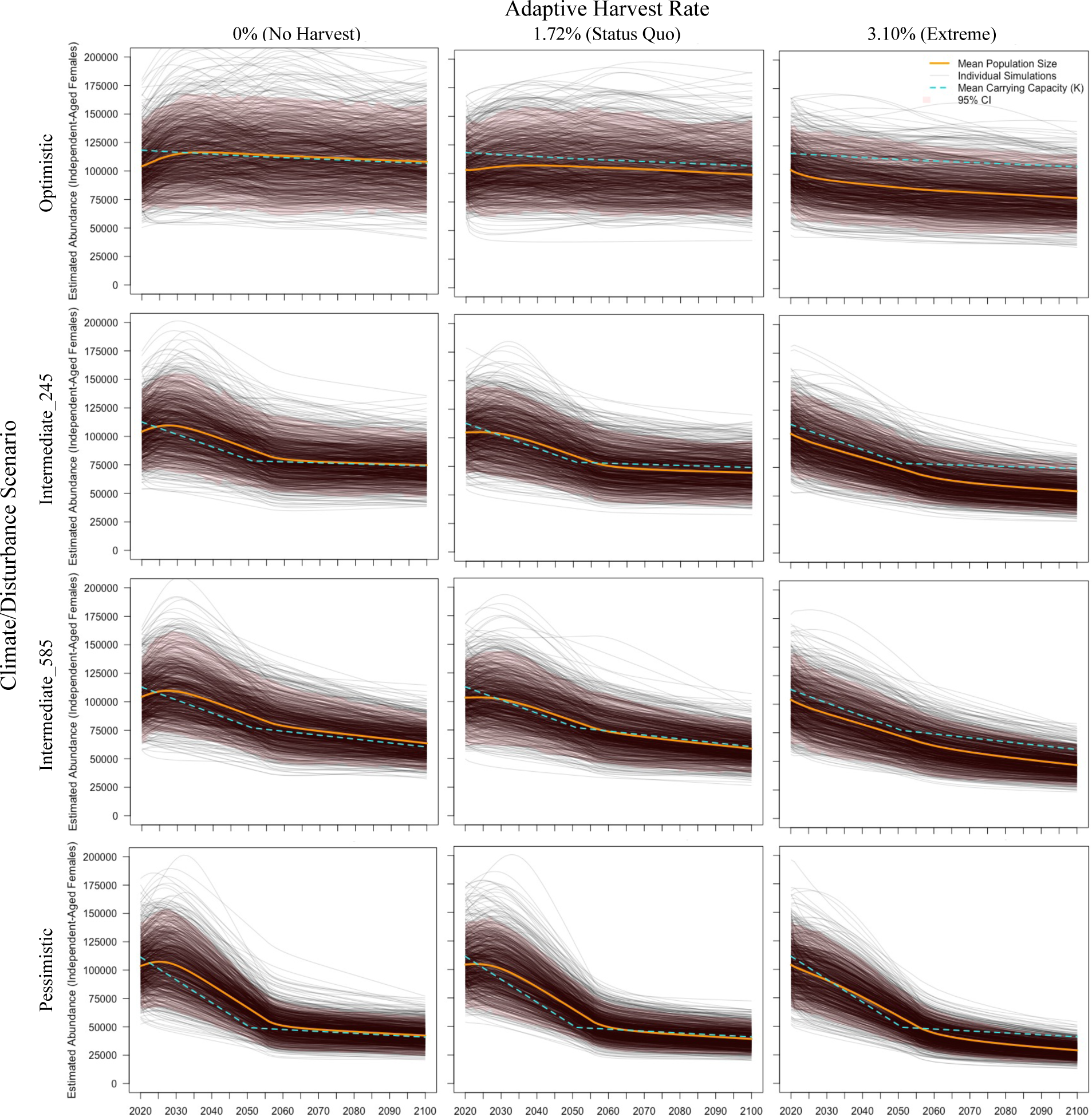
The effect of incorporating carrying capacity (*K*) and intrinsic rate of increase (*r_max_*) from four combined climate/disturbance scenarios (Optimistic; Intermediate_245; Intermediate_585; Pessimistic; details in Table 1) on modeled independent-aged female Pacific walrus population size, considering the three example harvest regimes: no harvest, status-quo harvest (1.72%), extreme harvest (3.1%) and a 10-year monitoring interval. In each panel, orange lines represent the mean estimated population size of independent-aged female Pacific walruses, gray lines are individual simulations (n=1000), cyan dashed lines represent mean carrying capacity estimates from each climate/disturbance scenario, and the red shaded region is a 95% credible interval of theta-logistic simulations. The Intermediate_245 row in Figure 6 corresponds to the top panels in figure 5. Associated management objective probabilities and harvest amounts for each model can be found in Table 1.

### Non-Adaptive vs Adaptive Harvest

Non-adaptive harvest scenarios (i.e., harvesting the same number of animals each year regardless of population size) resulted in higher probabilities of unsustainability (MO1) and quasi-extinction (MO2) than their adaptive harvest counterparts with similar initial harvest levels (Table 2; Tables S2-S5). As an example, we consider non-adaptive scenarios where 1,280 and 3,123 independent-aged females are harvested each year under the intermediate_245 and pessimistic climate/disturbance scenarios. These absolute numbers are 1.23% and 3.00% of our assumed initial independent-aged female population size (N_2020_ = 104,123; derived from Beatty et al., 2022). Whereas their adaptive counterparts (Fig. 5B-C; Fig. 6) all met both management objectives to the end of the 21^st^ century, the non-adaptive scenarios failed both MO1 and MO2 (Fig. 7, Table 2). In the Intermediate_245 non-adaptive status quo harvest scenario, simply superimposing the non-adaptive harvest on a slight background rate of population decline resulted in a probability of harvest unsustainability beyond the MO1 threshold by the end of the 21^st^ century (Fig. 7A). The Pessimistic non-adaptive high harvest scenario (3,123 females/year) failed both management objectives by the year 2054 and had a 100% probability of quasi-extinction by the year 2088 (Fig. 7D).

**Figure 7.**
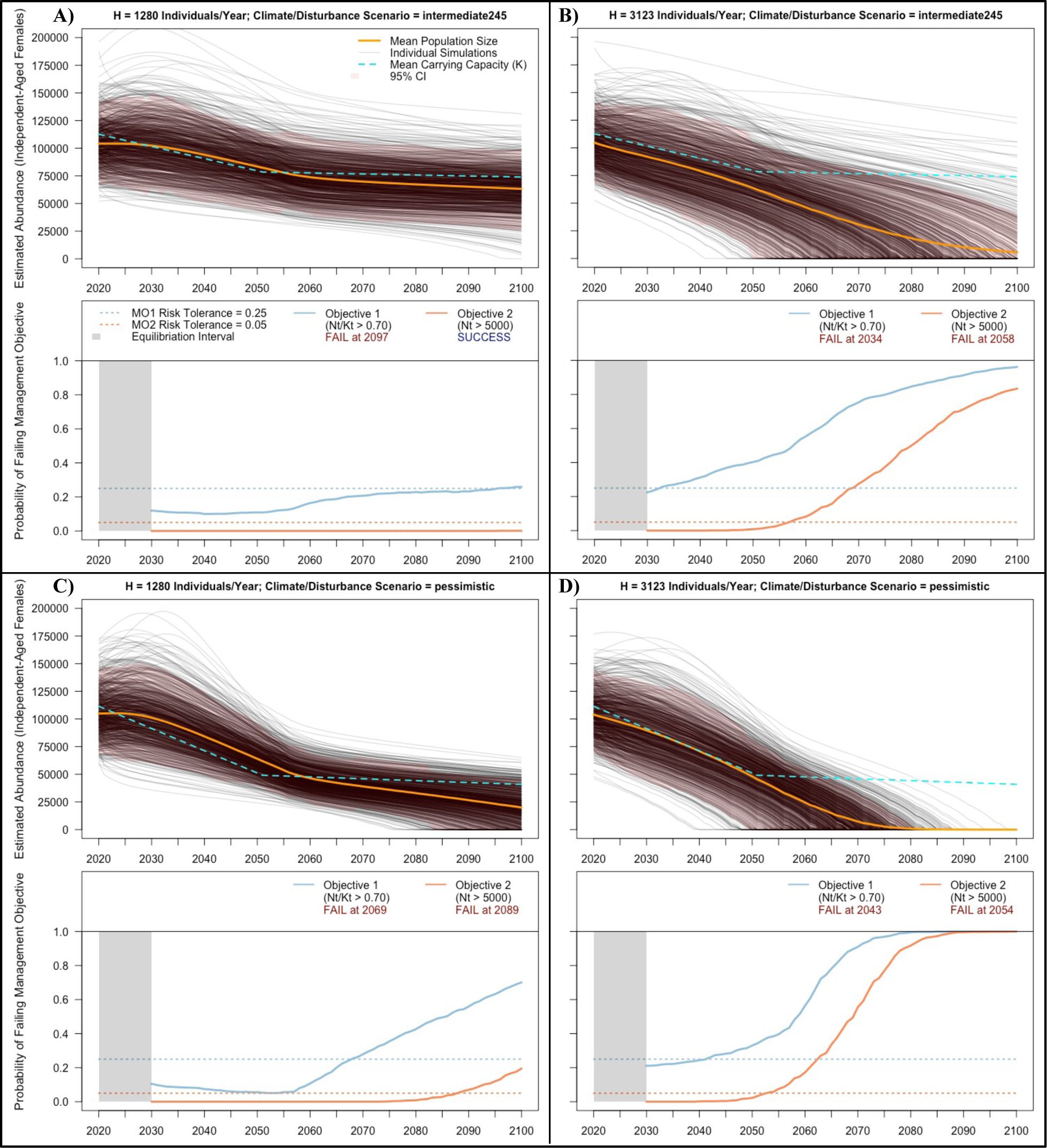
Example effects of non-adaptive harvest scenarios on independent-aged female Pacific walrus abundance. Top panels represent non-adaptive harvest scenarios with a constant annual harvest of 1280 individuals (A; equivalent to 1.23% of N_2020_) and 3123 individuals (B; equivalent to 3.00% of N_2020_) under an intermediate_245 climate/disturbance scenario. Bottom panels represent the same harvest scenarios under the pessimistic climate/disturbance scenario. In the top frame of each panel, orange lines represent the mean estimated population size of independent-aged female Pacific walruses, gray lines are individual simulations (n=1000), cyan dashed lines represent mean carrying capacity estimates from each climate/disturbance scenario, and the red shaded region is a 95% credible interval of theta-logistic simulations. Bottom frames represent associated probabilities of failing to meet risk tolerance thresholds for pre-defined management objectives (MO1 & MO2) at each timestep of the simulation.

### Monitoring Interval Sensitivity

The theta-logistic model was sensitive to the monitoring interval only at very high harvest rates (Fig. 8). Generally, decreasing the monitoring interval to 5 years resulted in higher population sizes at the end of the simulation, whereas increasing the interval to 15 years reduced the end population size and increased the quasi-extinction probability (Fig. 8; Table 2; Table S2-S5). However, there was very little difference in model output between the three monitoring intervals near status-quo rates of harvest (e.g., 1–1.5%, Tables S3–S5), regardless of climate/disturbance scenario. We observed significant differences (i.e., non-overlapping 95% credibility intervals at the end of the simulation) between the 5-year and 15-year monitoring intervals only when the simulated population was undergoing a severe decline (e.g., with 6% harvest; Fig. 8).

**Figure 8.**
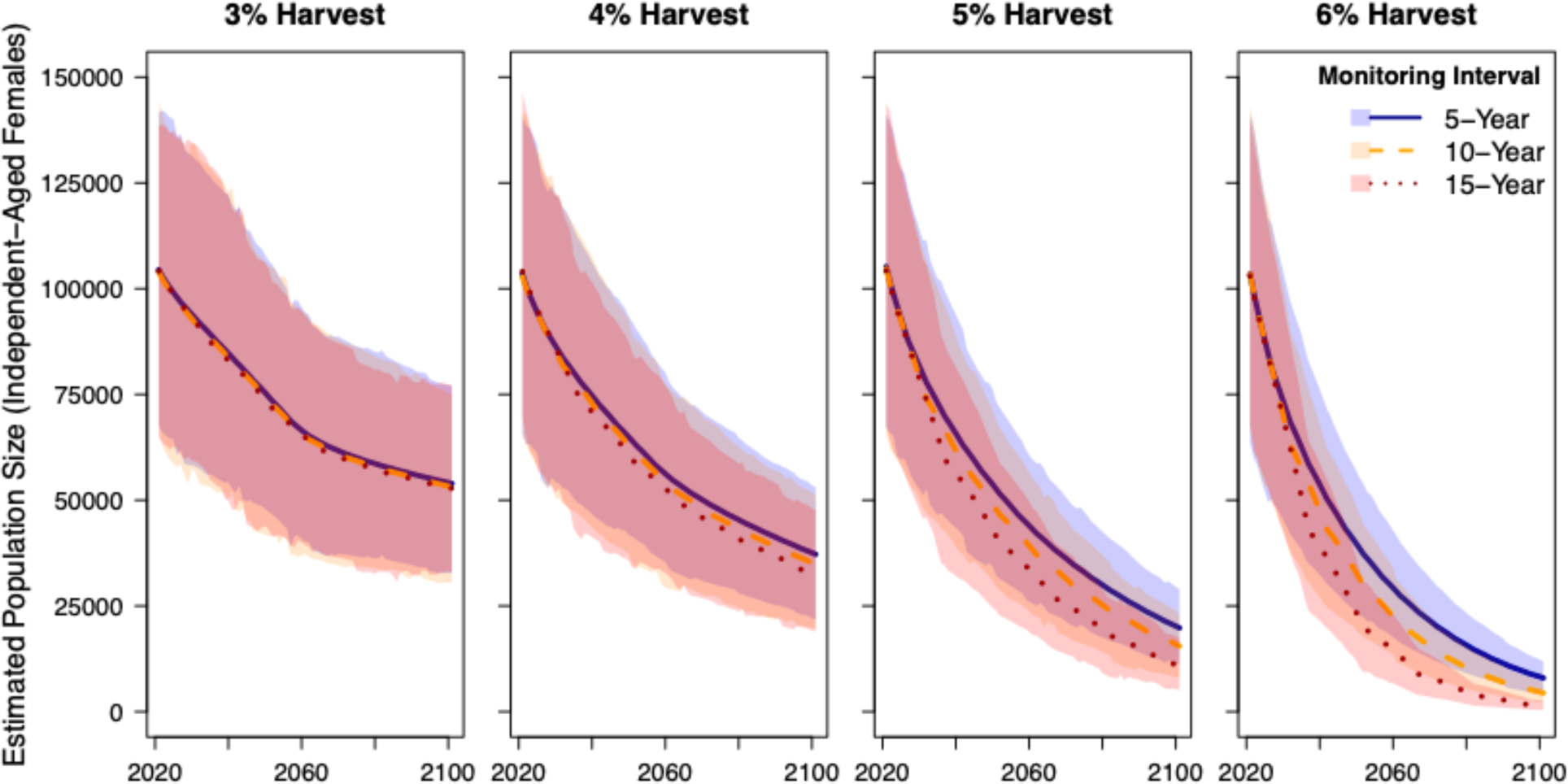
The effect of monitoring interval on projected population size in the theta-logistic model, considering four example harvest scenarios (3-6% harvest) in which the three monitoring interval options are compared. All models incorporate the intermediate_245 climate/disturbance scenario. In each panel, lines represent the mean estimated population size of independent-aged female Pacific walruses, and shaded regions represent a 95% credible interval of 1000 simulations for each model.

### Risk Tolerance Sensitivity

The theta-logistic model was sensitive to the risk tolerance level applied to MO1. We estimated the highest percentage-based harvest allowable to meet MO1 under the four combined climate/disturbance scenarios to the end of the 21^st^ century with low (15%), medium (25%), and high (35%) risk tolerance levels (Table S5). Across climate/disturbance scenarios, the maximum annual harvest rate of the independent-aged female Pacific walrus population that was sustainable was 1.26% when the risk tolerance level was 15%, 2.25% when the risk tolerance level was 25%, and 2.60% when the risk tolerance level was 35% (Table S5). An example of this risk tolerance sensitivity is demonstrated in Figure S3.

### Comparison Between Model Output & PBR

Overall, model output identified an adaptive annual harvest rate of 2.25% of the independent-aged female Pacific walrus population as the upper threshold to meet our criteria for sustainability (considering a 25% risk tolerance level) under all climate/disturbance scenarios considered to the end of the 21^st^ century. We calculated PBR for the independent-aged female Pacific walrus population (Supplemental Material 1), resulting in an estimate of 1313 individuals or 1.26% of the mean independent-aged female abundance estimate (*N_2020_* = 104,123). Thus, the harvest sustainability model indicated that a higher rate of harvest (an additional 0.99% of *N_2020_* or 1031 individuals) may be sustainable than that derived from the PBR equation. Nevertheless, if 1313 independent-aged females were harvested every year regardless of changes to population size, our simulations indicate that this constant quota may not be sustainable.

## DISCUSSION

The Pacific walrus population is projected to undergo a decline to the end of the 21^st^ century due to loss of sea ice habitat, but previous studies have been unable to estimate the magnitude and severity of that decline (MacCracken 2012; MacCracken et al., 2017). Recent studies have characterized various aspects of walrus space and energy usage in response to a changing climate (e.g., Fischbach et al., 2022; Udevitz et al., 2017) and anthropogenic disturbance (e.g., Taylor et al., 2023), which has allowed for the development of a framework to estimate future population-level parameters (i.e., the walrus PCoD model; Johnson et al., 2023). In the present study, we introduce a framework to couple those parameter estimates with different harvest scenarios to estimate levels of sustainable harvest and overall population outcomes.

We developed a quantitative harvest assessment for the Pacific walrus which showed potential population size changes in response to scenarios incorporating climate change, anthropogenic disturbance, and harvest. Although it is difficult to fully predict how walrus population size will respond to these factors, our scenarios provide a useful framework to assess harvest sustainability and quasi-extinction risk in a changing environment. The four climate/disturbance scenarios considered (Table 1) were designed to simulate a broad range of outcomes to the end of the 21^st^ century (Johnson et al., 2023). Generally, these scenarios indicated the walrus population will decline over time even in the absence of harvest (e.g., Fig. 6). However, sustainable adaptive harvest scenarios (i.e., which maintain *N/K* > 0.7; MO1) are still possible under these population declines. Specifically, our models indicated an annual adaptive harvest rate of 2.25% of the independent-aged female population (which is higher than the range of contemporary harvest levels) is near the upper threshold of sustainability across climate scenarios to the end of the 21^st^ century.

Harvest sustainability is a primary goal of hunters and wildlife managers alike, but there is substantial ambiguity among the scientific community regarding how sustainability should be defined and measured (e.g., Weinbaum et al., 2014). The PBR equation is commonly used for marine mammal stocks in the United States (Wade 1998) and has been used as a standard of sustainability in previous Stock Assessment Reports for the Pacific walrus (USFWS, 2023).

Although it might be useful to have a standardized metric for harvest sustainability, PBR is a simple equation (Punt et al., 2020) that likely does not adequately encompass the complexities associated with harvested populations undergoing widespread changes in habitat (Robards et al., 2009). The PBR method to calculate mortality limits for marine mammals reflects built-in management objectives and risk tolerances that seek to minimize unwanted mortality associated with fisheries bycatch. However, in some cases PBR is considerably lower than the level of removals that would be considered sustainable for subsistence harvest (e.g., polar bears in the Chukchi Sea; Regehr et al. 2021). The theta-logistic modeling approach we applied in this study is a way to account for those complexities. In addition, PBR is traditionally calculated based on the number of individuals in all sex-age classes in the population. This can be problematic for species that are polygynous, such as the Pacific walrus, and still have relatively large population sizes, as these factors combine to make adult males demographically unimportant (e.g., Taylor et al., 2018). Our female-based theta-logistic modeling also addresses this issue. We calculated PBR for the independent-aged female component of the Pacific walrus population to compare against our estimates for the maximum sustainable yield from the theta-logistic model (Supplemental Material 1). The PBR estimate (1,313 independent-aged female Pacific walruses; 1.26% of the assumed independent-aged female population size in 2020) was below the highest percentage-based harvest that met our management objective for sustainability in the theta-logistic model to the end of the 21^st^ century (between 2% and 2.5%; Fig. 3). The relatively low proportion of animals that would be harvested based on the PBR compared to the theta-logistic model largely reflects a conservative recovery factor value (F_R_ = 0.5, Supplemental Material 1) that allows room for error due to known biases and uncertainties associated with human-caused mortality rates and future population trends. In fact, without this recovery factor, the PBR equation would indicate that 2,626 independent-aged female walruses could have been sustainably harvested in 2020 (a rate of 2.5%), which, according to the theta-logistic model would not be sustainable to the end of the century. Notably, a constant harvest of 1,313 independent-aged females may not be sustainable (depending on the climate/disturbance scenario), and constant harvest without the recovery factor (2,626) would not be sustainable and would lead to quasi-extinction before the end of the 21^st^ century. This underscores the importance of adaptive harvest rates, wherein the recommended sustainable harvest rate is calculated as a percentage of the population and reassessed periodically as new population estimates become available (e.g., Fig. 1).

This type of adaptive harvest management assumes a harvest co-management framework exists for the Pacific walrus population (and continues to exist to the end of the 21^st^ century) that can obtain regular abundance estimates (at 5, 10, or 15-year intervals), monitor harvest, and adhere to sustainable harvest estimates. This is consistent with the 10-year population reassessment interval suggested by MacCracken et al. (2017), but could change if resources are significantly reallocated, or as methods for abundance estimates continue to develop (e.g., genetic mark-recapture, Beatty et al., 2022). Harvest monitoring programs are active in both the United States and Russia and report high levels of hunter compliance (∼95% for Alaska, ∼80% for Russia; MacCracken et al., 2017; Smirnov et al., 2002). Annual harvest quotas are set by the Russian government for Chukotkan communities (Kryukova 2019), whereas Alaskan walrus-harvesting communities operate without a firm quota. Some Alaskan communities have established Marine Mammal Advisory Committees which have implemented ordinances to provide “bag limits” in some instances in the past which stipulate that salvage should not be wasteful (limiting the number of animals that can be killed in any single foray; MacCracken et al., 2017). Ultimately, implementing an adaptive harvest for a changing Pacific walrus population will require continued close coordination and co-management between wildlife agencies in the United States and Russia and with subsistence walrus-hunting communities.

Pacific walrus population abundance estimates have historically been infrequent, costly and logistically challenging to acquire coupled with low precision (e.g., Table S1). The most recent population estimate used modern genetic mark-recapture techniques (Beatty et al., 2022), which resulted in a more precise estimate than has been possible historically. However, obtaining the data required for this approach is challenging because of its high costs and complex logistics (Beatty et al. 2022). We introduced the “monitoring interval” parameter (5, 10, or 15 years) into the theta-logistic adaptive harvest scenarios to assess the importance of the frequency of population abundance estimates on simulated population outcomes (Fig. 8). Model output was very similar between these three monitoring intervals, and only exhibited significant differences when the population was undergoing severe declines associated with very high harvest levels (e.g., 6% of the population, Fig. 8). Apart from harvest, any factor contributing to a severe and rapid population decline (e.g., some catastrophic climate-related event more severe than our most pessimistic climate/disturbance scenario) would also necessitate a reconsideration of the monitoring interval. In such an event, anecdotal evidence of a rapidly declining population (e.g., reports of massive die-offs at haulouts; concerns from subsistence hunters) may warrant reconsideration of the monitoring interval or a temporary adjustment to harvest percentages or risk tolerance values. However, in most realistic cases, the cost may outweigh the benefit of increasing the frequency of population estimates with regards to harvest management.

Along with adaptive harvest scenarios, we analyzed a suite of “non-adaptive” scenarios where a constant number of individuals were harvested annually throughout the course of the simulation, which could represent cases where a minimum number of animals must be harvested to meet community needs, irrespective of the status of the population. Generally, these non-adaptive scenarios resulted in higher probabilities of unsustainability or quasi-extinction compared to adaptive harvest rates that scale to a changing population size. As visualized in Figure 7B, when a status-quo harvest scenario is combined with even the slight decline in population size associated with an Intermediate_245 climate/disturbance scenario, the population is driven below its sustainability threshold (MO1). Realistically, non-adaptive scenarios may become unlikely as the population declines over time—because the availability of walrus to hunters could decrease with decreasing abundance, which could regulate the harvest in a density-dependent manner before the population nears its quasi-extinction threshold. Overall, our results stress the importance of adaptive management plans for populations that are undergoing rapid environmental change.

The management objectives in this study were developed as a collaborative effort between wildlife managers and IK holders from subsistence walrus hunting communities. Our primary criterion for harvest sustainability was Management Objective 1: maintaining population size above MNPL—a metric recommended for harvested populations in changing environments (Regehr et al., 2021a; USFWS 2016). Management Objective 2 (maintaining the population above a quasi-extinction threshold) represents a worst-case scenario and has further implications for the species’ status should it be exceeded (e.g., listing under the Endangered Species Act). In the present study we apply a placeholder value of 5,000 independent-aged female Pacific walruses as the point at which deleterious small-population effects may occur, but this value could be updated as more information becomes available on walrus population genetics. Additional management objectives (e.g., maintaining population abundance over a certain threshold; requiring set minimum harvest levels to meet specific community needs) could easily be incorporated into future versions of the harvest assessment framework.

Risk tolerance thresholds are an important component of this analysis, and the selection of such thresholds is a decision that warrants careful discussion and coordination between managers and stakeholders. Although we summarize results considering a medium (25%) risk tolerance threshold for our sustainability management objective (MO1), we demonstrated that alternative risk tolerance levels will yield different interpretations of sustainable harvest strategies (Table S5; Fig. S3). The flexibility of the modeling framework in this regard allows managers and subsistence users to quantitatively consider the risk associated with various harvest strategies under different climate/disturbance scenarios to ultimately consider the harvest strategy that will best meet community needs while promoting harvest sustainability into the future.

The theta-logistic modeling approach we apply in this study is a useful framework to conceptualize the sustainability of harvest under a wide array of scenarios, but it is ultimately a simplification of complex natural processes and population dynamics that relies on a series of assumptions. Most notably, it focuses on the independent-aged female component of the population (following Regehr et al., 2021) because females are the most important reproductive drivers of polygynous marine mammal populations (e.g., Laidre et al., 2008) and because estimates of *K* and *r_max_* from the PCoD model were derived specifically for independent-aged females (Johnson et al., 2023). Harvest assessments incorporating more complex age-structured matrix models have been developed for other species (e.g., Regehr et al., 2017; Regehr et al., 2021b) but require a higher quality of sex and age-specific harvest and population data than is currently available for the Pacific walrus population. Thus, we were unable to evaluate scenarios that assess different sex and age compositions of harvest, and instead must assume that contemporary harvest demographics (e.g., Fig. S1) remain consistent in the future. An increase in the quality of demographic harvest data may allow for the incorporation of age- and sex-structured matrix models in future iterations of this harvest assessment.

## MANAGEMENT IMPLICATIONS

Wildlife species that are undergoing rapid habitat loss associated with climate change and are also important subsistence resources to local communities represent a unique ecological and socio-economic challenge, requiring an effective and adaptive management strategy to promote sustainable harvest. The harvest assessment framework we developed in the present study provides evidence that sustainable harvests are possible for a declining Pacific walrus population, but that harvest levels would need to adapt if and when the population size changes, which is expected based on projected conditions for walruses. Climate-driven habitat loss is affecting wildlife populations on a global scale and using such frameworks to construct and assess adaptive management scenarios will become increasingly important for wildlife managers and subsistence hunters.

## ACKNOWLEDGEMENTS

We are extremely thankful to members of the Pacific Walrus Harvest Model Steering Committee and the Eskimo Walrus Commission for their support and discussion in the development of this modeling framework: Charles Brower, Vera Metcalf, Jacob Martin, Bryan Rookuk Jr., and Enoch Oktollik. Jen Cate and Aaron Crist provided useful input on model development and William Beatty and Martin Robards provided thoughtful reviews on previous versions of this manuscript. The findings and conclusions in this article are those of the author(s) and do not necessarily represent the views of the US Fish and Wildlife Service. This product paper has been peer reviewed and approved for publication consistent with USGS Fundamental Science Practices (https://pubs.usgs.gov/circ/1367/). Any use of trade, firm or product names is for descriptive purposes only and does not imply endorsement by the U.S. Government.

## Works Cited

Beatty WS, PR Lemons, JP Everett, CJ Lewis, RL Taylor, RJ Lynn, SA Sethi, L Quakenbush, JJ Citta, ML Kissling, N Kryukova & JK Wenburg. 2022. Estimating Pacific walrus abundance and survival with multievent mark-recapture models. Marine Ecology Progress Series 697: 167–182. doi: 10.3354/meps14131

Cortes E. 2016. Perspectives on the intrinsic rate of population growth. Methods in Ecology and Evolution 7: 1136–1145. doi: 10.1111/2041-210X.12592

Fay FH. 1982. Ecology and biology of the Pacific walrus, Odobenus Rosmarus divergens illiger. USFWS North American Fauna (74).

Fay, FH, BP Kelly & JL Sease. 1989. Managing the exploitation of Pacific walruses: a tragedy of delayed response and poor communication. Marine Mammal Science 5:1–16.

Fay FH, JJ Burns, SW Stoker, JS Grundy. 1994. The struck-and-lost factor in Alaskan walrus harvests, 1952-1972. Arctic 47(4): 368–373.

Fischbach AS, DH Monson & CV Jay. 2009. Enumeration of Pacific walrus carcasses on beaches of the Chukchi Sea in Alaska following a mortality event, September 2009. U.S. Department of the Interior, U.S. Geological Survey Open-File Report, 2009–129, Reston, VA.

Fischbach AS, RL Taylor & CV Jay. 2022. Regional walrus abundance estimate in the United States Chukchi Sea in autumn. Journal of Wildlife Management 82:e22256. doi: 10.1002/jwmg.22256.

Fox-Kemper, B, HT Hewitt, C Xiao, G Aðalgeirsdóttir, SS Drijfhout, TL Edwards, NR Golledge, M Hemer, RE Kopp, G Krinner, A Mix, D Notz, S Nowicki, IS Nurhati, L Ruiz, J-B Sallée, ABA Slangen, & Y Yu. 2021. Ocean, Cryosphere and Sea Level Change. In Climate Change 2021: The Physical Science Basis. Contribution of Working Group I to the Sixth Assessment Report of the Intergovernmental Panel on Climate Change. [Masson-Delmotte, V, P Zhai, A Pirani, SL Connors, C Péan, S Berger, N Caud, Y Chen, L Goldfarb, MI Gomis, M Huang, K Leitzell, E Lonnoy, JBR Matthews, TK Maycock, T Waterfield, O Yelekçi, R Yu, and B Zhou (eds.)]. Cambridge University Press, Cambridge, United Kingdom and New York, NY, USA, pp. 1211–1362, doi:10.1017/9781009157896.011.

Gadamus L & J Raymond-Yakoubian. 2015. A Bering Strait Indigenous framework for resource management: respectful seal and walrus hunting. Arctic Anthropology 52(2): 87–101.

Garlich-Miller JL, LT Quakenbush & JF Bromhagin. 2006. Trends in age structure and productivity of Pacific walruses harvested in the Bering Strait region of Alaska, 1952-2002. Marine Mammal Science 22(4): 880–896. doi: 10.1111/j.1748-7692.2006.00081.x.

Garlich-Miller JL & DM Burn. 2009. Estimating the harvest of Pacific walrus, Odobenus rosmarus divergens, in Alaska. Fishery Bulletin – National Oceanic and Atmospheric Administration 97(4): 1043–2046.

Hovelsrud GK, M Mckenna & HP Huntington. 2008. Marine mammal harvests and other interactions with humans. Ecological Applications 18:S135–S147.

Inuit Circumpolar Council-Alaska. 2015. Alaskan Inuit Food Security Conceptual Framework: how to assess the Arctic from an Inuit perspective. Technical Report. https://iccalaska.org/wp-icc/wp-content/uploads/2016/05/Food-Security-Full-Technical-Report.pdf.

Johnson FA, M Alhainen, AD Fox, J Madsen & M Guillemain. 2018. Making do with less: must sparse data preclude informed harvest strategies for European waterbirds? Ecological Applications 28: 427–441. doi: 10.1002/eap.1659.

Johnson DL, JM Eisaguirre, RL Taylor & JL Garlich-Miller. 2023. Assessing the population consequences of disturbance and climate change for the Pacific walrus (Pre-print). bioRxiv doi: 10.1101/2023.10.12.562073.

Kryukova, NV. 2019. Legal regulation of the traditional (native) Pacific walrus harvest in Russia. Artika I Sever [Arctic and the North] 36: 24–41. doi: 10.17238/issn2221-2698.2019.36.24.

Kryukova, N.V. 2019. Monitoring the harvest of Pacific walrus in Russia. Kamchatka Branch of the Pacific Geographical Institute FEB RAS. 16 p. doi: 10.17238/issn2221-2698.2019.36.24.

Laidre KL, I Stirling, LF Lowry, O Wiig, MP Heide-Jorgensen & SH Ferguson. 2008. Quantifying the sensitivity of Arctic marine mammals to climate-induced habitat change. Ecological Applications 18(2): S97–S125. doi: 10.1890/06-0546.1.

Lubow BC, BL Smith. 2010. Population dynamics of the Jackson elk herd. Journal of Wildlife Management 68(4): 810–829. doi: 10.2193/0022-541X(2004)068[0810:PDOTJE]2.0.CO;2.

MacCracken JG. 2012. Pacific Walrus and climate change: observations and predictions. Ecology and Evolution 2(8): 2072–2090. doi: 10.1002/ece3.317.

MacCracken JG, WS Beatty, JL Garlich-Miller, ML Kissling, & JA Snyder. Final species status assessment or the Pacific walrus (Odobenus rosmarus divergens), May 2017 (Version 1.0). United States Fish and Wildlife Service, Marine Mammals Management. Technical report.

Metcalf V & M Robards. 2008. Sustaining a healthy human-walrus relationship in a dynamic environment: challenges for comanagement. Ecological Applications 18(2): S148–S156. doi: 10.1890/06-0642.1.

Metcalf V, HP Huntington, JL Garlich-Miller & DL Johnson. 2023. Walruses and the future: incorporating Indigenous Knowledge into Pacific walrus population and harvest sustainability models (interim title). [Workshop Report, in press]. Nome, AK, August 22-24.

Novachek MJ, EE Cleland. 2001. The current biodiversity extinction event: Scenarios for mitigation and recovery. Proceedings of the National Academy of Sciences 19(10): 5466–5470. doi: 10.1073/pnas.091093698.

Nilsen EB, O Strand. 2018. Integrating data from multiple sources for insights into demographic processes: Simulation studies and proof of concept for hierarchical change-in-ratio models. PLoS ONE 13(3): e094556. doi: 10.1371/journal.pone.0194566.

National Marine Fisheries Service (NMFS). 2016. Guidelines for preparing stock assessment reports pursuant to the 1994 amendments to the Marine Mammal Protection Act. 23 p. Available online: https://www.fisheries.noaa.gov/national/marine-mammal-protection/guidelines-assessing-marine-mammal-stocks. Accessed June 2022.

Overland J, E Dunlea, JE Box, R Corell, M Forsius, V Kattsov, M Skovgard Olsen, J Pawlak, L Reiersen & M Wang. 2019. The urgency of Arctic change. Polar Science 21 (2019) 6–13. doi: 10.1016/j.polar.2018.11.008.

Post E, Forchhammer MC, Bret-Harte MS, Callaghan TV, Christensen TR, Elberling B, Fox AD, Gilg O, Hik DS, Høye TT, Ims RA, Jeppesen E, Klein DR, Madsen J, McGuire AD, Rysgaard S, Schindler DE, Stirling I, Tamstorf MP, Tyler NJC, van der Wal R, Welker J, Wookey PA, Schmidt NM, Aastrup P. 2009. Ecological dynamics across the Arctic associated with recent climate change. Science 325: 1355–1358. doi: 10.1126/science.1173113.

Punt AE, M Siple, TB Francis, PS Hammond, D Heinemann, KJ Long, JE More, M Sepulveda, RR Reeves, GM Sigurdsson, G Vikingsson, PR Wade, R Williams & AN Zerbini. Robustness of potential biological removal to monitoring, environmental, and management uncertainties. ICES Journal of Marine Science 77: 2491–2507. doi: 10.1093/icesjms/fsaa096.

Ragen, T. 1995. Maximum net productivity level estimation for the Northern fur seal (*Callorhinus ursinus*) population of St. Paul Island, Alaska. Marine Mammal Science 11(3):275–300.

Reed DH, JJ O’Grady, BW Brook, JD Ballou & Richard Frankham. 2003. Estimates of minimum viable population sizes for vertebrates and factors influencing those estimates. Biological Conservation 113: 23–34. doi: 10.1016/S0006-3207(02)00346-4.

Regehr EV, RR Wilson, KD Rode, MC Runge & HL Stern. 2017. Harvesting wildlife affected by climate change: a modelling and management approach for polar bears. Journal of Applied Ecology 54: 1534–1543. doi: 10.1111/1365-2664.12864.

Regehr EV, M Dyck, S Iverson, DS Lee, NJ Lunn, JM Northrup, M Richer, G Szor & MC Runge. 2021a. Incorporating climate change in a harvest risk assessment for polar bears *Ursus maritmus* in Southern Hudson Bay. Biological Conservation 258 (2021) 109128. doi: 10.1016/j.biocon.2021.109128.

Regehr EV, MC Runge, AV Duyke, RR Wilson, L Polasek KD Rode, NJ Hostetter & SJ Converse. 2021b. Demographic risk assessment for a harvested species threatened by climate change: polar bears in the Chukchi Sea. Ecological Applications 31(8): e02461. doi: 10.1002/eap.2461.

Robards MD & JL Joly. 2008. Interpretation of “wasteful manner” within the Marine Mammal Protection Act and its role in management of the Pacific walrus. Ocean and Coastal Law Journal 13(2): 171–232.

Robards MD, JJ Burns, CL Meek & A Watson. 2009. Limitations of an optimum sustainable population or potential biological removal approach for conserving marine mammals: Pacific walrus case study. Journal of Environmental Management 91: 57–66. doi: 10.1016/j.jenvman.2009.08.016.

Smirnov G, V Rinteimit, M Agnakisyak, M Likovka. 2002. Walrus harvest monitoring on Chukotka in 2001. USFWS technical report, April 2002. Chukotka Branch of the Pacific Fisheries Research Center.

Taylor RL & MS Udevitz. 2015. Demography of the Pacific walrus (*Odobenus rosmarus divergens*): 1974-2006. Marine Mammal Science 31(1): 231–254. doi: 10.1111/mms.12156.

Taylor RL, MS Udevitz, CV Jay, JJ Citta, LT Quakenbush, PR Lemons, & JA Snyder. 2018. Demography of the Pacific walrus (*Odobenus rosmarus divergens*) in a changing Arctic. Marine Mammal Science 34(1): 54–86. doi: 10.1111/mms.12434.

Taylor RL, CV Jay, WS Beatty, AS Fischbach, LT Quakenbush & JA Crawford. Exploring effects of vessels on walrus behaviors using telemetry, automatic identification system data and matching. Ecosphere 14:e4433. doi: 10.1002/ecs2.4433.

Udevitz MS, RL Taylor, JL Garlich-Miller, LT Quakenbush & JA Snyder. 2012. Potential population-level effects of increased haulout-related mortality of Pacific walrus calves. Polar Biology 36: 291–298. doi: 10.1007/s00300-012-1259-3.

Udevitz MS, CV Jay, RL Taylor, AS Fischbach, WS Beatty & SR Noren. 2017. Forecasting consequences of changing sea ice availability for Pacific walruses. Ecosphere 8(11): e02014.

USFWS – United States Fish and Wildlife Service, 2016. Polar Bear (Ursus maritimus) Conservation Management Plan, Final. US Fish and Wildlife Service, Region 7, Anchorage, Alaska. Available at. https://www.fws.gov/alaska/pages/what-we-do/marine-mammals/polar-bear-program/Plan.

USFWS (U.S. Fish and Wildlife Service). 2023. Pacific walrus (*Odobenus rosmarus divergens*): Alaska Stock. U.S. Department of the Interior, U.S. Fish and Wildlife Service, Marine Mammals Management, Anchorage, AK.

Wade, P.R., 1998. Calculating limits to the allowable human–caused mortality of cetaceans and pinnipeds. Marine Mammal Science 14:1–37. doi: 10.1111/j.1748-7692.1998.tb00688.x

Weinbaum KZ, JS Brashares, CD Golden & WM Getz. 2013. Searching for sustainability: are assessments of wildlife harvests behind the times? Ecol Lett. 16(1): 99–111. doi:10.1111/ele.12008.

Williams PJ & MB Hooten. 2016. Combining statistical inference and decisions in ecology. Ecological Applications 26(6): 1930–1942. doi: 10.1890/15-1593.1.

