## Supplemental Material for "Assessing the Sustainability of Pacific Walrus Harvest in a Changing Environment"

**For**

**Supplemental Material 1.**

Calculation of Potential Biological Removal (PBR) for the Independent-aged female Component of the Pacific Walrus Population

According to the Marine Mammal Protection Act, the potential biological removal (PBR) of a marine mammal stock is defined as:

*“The maximum number of animals, not including natural mortalities that may be removed from a marine mammal stock while allowing that stock to reach or maintain its optimal sustainable population*. *The PBR is the product of the following factors: (A) the minimum population estimate N_MIN_, (B) one-half the maximum net productivity rate (0.5 R_MAX_), and (C) a recovery factor (F_R_) ranging between 0.1 and 1.0* (MMPA §3(20))”

This can be expressed as the following equation:

*PBR* = *N_MIN_* * 0.5 (*R_MAX_*) * *F_R_*	(Equation 1)

According to the official NMFS guidelines (NMFS 2016), *N_MIN_* for the PBR equation should approximate the 20^th^ quantile of a log-normal distribution used to describe the estimated population size. The most recent abundance estimate of the Pacific walrus population (Beatty et al., 2022) was used to derive a distribution of abundance for independent-aged females (95% CrI = 65,827–142,418). Thus, for the *N_MIN_* value for the PBR equation, we simply calculated the 20^th^ quantile of the posterior distribution of the independent-aged female abundance estimate (*N_MIN_* = 87,526). We applied values of *R_MAX_* (0.06) and *F_R_* (0.5) that were used in the most recent Pacific walrus stock assessment (Garlich-Miller et al., 2022). This resulted in the following estimate of PBR:

PBR = 87,526 * 0.5 (0.06) * 0.5 = 1312.89	(Equation 2)

For comparison to theta-logistic model results, we can scale PBR proportionally to the assumed initial population size for independent-aged females we used in our simulations (N_2020_ = 104,123), which equates to an annual percentage-based harvest of 1.26% of the population.

**Supplemental Tables:**

Table S1. Pacific walrus abundance estimates (1975–2015) with 95% confidence or credible intervals. Estimates are derived from one of three methods: Aerial survey/terrestrial haulout survey (AS/THS); Aerial survey/infra-red (AS/IR); and genetic mark-recapture (GMR).

| Year | Total Abundance  Mean | Total Abundance  95% CI/CrI | Type | Source |
| --- | --- | --- | --- | --- |
| 1975^1,2^ | 199,738 | 112,000–330,000 | AS/THS | Estes & Gol’tsev (1984) |
| 1980^1,2^ | 254,890 | 184,000–344,000 | AS/THS | Johnson et al. (1982) & Fedoseev (1981) |
| 1985^1,2^ | 242,882 | 125,000–427,000 | AS/THS | Gilbert (1989); Fedoseev & Razlivalov (1986) |
| 1990^2^ | 201,039 | 88,000–397,000 | AS/THS | Gilbert et al., (1992) |
| 2006 | 129,000 | 55,000–507,000 | AS/IR | Speckman et al., (2011) |
| 2015 | 257,193 | 171,138–366,366 | GMR | Beatty et al., (2022) |

^1^ On-ice components of these abundance estimates were subsequently adjusted by Udevitz et al., (2001).

^2^ Confidence intervals were retroactively estimated by Taylor & Udevitz (2015).

Table S2. Summary of theta-logistic model output considering 44 harvest scenarios under the Optimistic climate/disturbance scenario. H_type (harvest type) is either A (adaptive scenarios) or N-A (non-adaptive scenarios); M_int is the monitoring interval; H is the percentage of the independent-aged female Pacific walrus population that is harvested annually in each scenario; A_t_ is the absolute harvest under each scenario; N_t_ is the mean estimated population size at each timestep of the simulation (i.e., the years 2050 and 2100); MO1 and MO2 are the probabilities of meeting Management Objective 1 (sustainability) and Management Objective 2 (avoiding quasi-extinction). Simulations which fail either management objective on at least one criterion according to the pre-defined risk tolerance thresholds (25% for MO1, 5% for MO2) are bolded.

| **H_type** | **M_int** | **H** | **A_2020_** | **N_2050_** | **A_2050_** | **MO1_2050_** | **MO2_2050_** | **N_2100_** | **A_2100_** | **MO1_2100_** | **MO2_2100_** |
| --- | --- | --- | --- | --- | --- | --- | --- | --- | --- | --- | --- |
| - | - | 0.00% | 0 | 113734 | 0 | 0.108 | 0.000 | 107559 | 0 | 0.114 | 0.000 |
| A | 5 | 0.50% | 520 | 111321 | 556 | 0.11 | 0.000 | 106092 | 530 | 0.124 | 0.000 |
| A | 5 | 1.00% | 1041 | 108032 | 1080 | 0.136 | 0.000 | 102424 | 1024 | 0.124 | 0.000 |
| A | 5 | 1.50% | 1561 | 104402 | 1566 | 0.142 | 0.000 | 98608 | 1479 | 0.160 | 0.000 |
| A | 5 | 2.00% | 2082 | 99605 | 1992 | 0.216 | 0.000 | 92904 | 1858 | 0.227 | 0.000 |
| A | 5 | **2.50%** | 2603 | 94844 | 2371 | **0.256** | 0.000 | 88372 | 2209 | **0.278** | 0.000 |
| A | 5 | **3.00%** | 3123 | 87695 | 2630 | **0.341** | 0.000 | 79414 | 2382 | **0.423** | 0.000 |
| A | 5 | **3.50%** | 3644 | 80807 | 2828 | **0.479** | 0.000 | 68516 | 2398 | **0.659** | 0.000 |
| A | 5 | **4.00%** | 4164 | 71481 | 2859 | **0.65** | 0.000 | 53358 | 2134 | **0.912** | 0.000 |
| A | 5 | **4.50%** | 4685 | 62827 | 2827 | **0.815** | 0.000 | 38492 | 1732 | **0.994** | 0.000 |
| A | 5 | **5.00%** | 5206 | 53021 | 2651 | **0.933** | 0.000 | 25442 | 1272 | **1.000** | 0.000 |
| A | 5 | **5.50%** | 5726 | 44698 | 2458 | **0.987** | 0.000 | 16359 | 899 | **1.000** | 0.000 |
| A | 5 | **6.00%** | 6247 | 37190 | 2231 | **1.000** | 0.000 | 10209 | 612 | **1.000** | 0.004 |
| A | 10 | 0.50% | 520 | 111230 | 560 | 0.126 | 0.000 | 104770 | 523 | 0.111 | 0.000 |
| A | 10 | 1.00% | 1041 | 108626 | 1094 | 0.125 | 0.000 | 102087 | 1020 | 0.124 | 0.000 |
| A | 10 | 1.50% | 1561 | 104568 | 1581 | 0.152 | 0.000 | 98108 | 1471 | 0.170 | 0.000 |
| A | 10 | 2.00% | 2082 | 99614 | 2008 | 0.224 | 0.000 | 93638 | 1872 | 0.221 | 0.000 |
| A | 10 | **2.50%** | 2603 | 94586 | 2391 | **0.266** | 0.000 | 87695 | 2192 | **0.288** | 0.000 |
| A | 10 | **3.00%** | 3123 | 87897 | 2676 | **0.342** | 0.000 | 79382 | 2381 | **0.430** | 0.000 |
| A | 10 | **3.50%** | 3644 | 78912 | 2829 | **0.517** | 0.000 | 66852 | 2339 | **0.681** | 0.000 |
| A | 10 | **4.00%** | 4164 | 69110 | 2874 | **0.716** | 0.000 | 50484 | 2019 | **0.942** | 0.000 |
| A | 10 | **4.50%** | 4685 | 58903 | 2817 | **0.873** | 0.000 | 33983 | 1529 | **0.998** | 0.000 |
| A | 10 | **5.00%** | 5206 | 48996 | 2683 | **0.964** | 0.000 | 20825 | 1041 | **1.000** | 0.000 |
| A | 10 | **5.50%** | 5726 | 39031 | 2438 | **0.997** | 0.000 | 11647 | 640 | **1.000** | 0.001 |
| A | 10 | **6.00%** | 6247 | 29386 | 2089 | **1.000** | 0.000 | 5877 | 352 | **1.000** | 0.275 |
| **H_type** | **M_int** | **H** | **A_2020_** | **N_2050_** | **A_2050_** | **MO1_2050_** | **MO2_2050_** | **N_2100_** | **A_2100_** | **MO1_2100_** | **MO2_2100_** |
| A | 15 | 0.50% | 520 | 111090 | 558 | 0.113 | 0.000 | 105027 | 528 | 0.104 | 0.000 |
| A | 15 | 1.00% | 1041 | 107918 | 1086 | 0.135 | 0.000 | 101146 | 1019 | 0.145 | 0.000 |
| A | 15 | 1.50% | 1561 | 103880 | 1570 | 0.178 | 0.000 | 97913 | 1478 | 0.183 | 0.000 |
| A | 15 | 2.00% | 2082 | 100935 | 2035 | 0.199 | 0.000 | 94918 | 1911 | 0.203 | 0.000 |
| A | 15 | **2.50%** | 2603 | 93741 | 2366 | **0.294** | 0.000 | 87620 | 2208 | **0.297** | 0.000 |
| A | 15 | **3.00%** | 3123 | 87519 | 2663 | **0.363** | 0.000 | 79607 | 2414 | **0.437** | 0.000 |
| A | 15 | **3.50%** | 3644 | 78979 | 2830 | **0.512** | 0.000 | 66946 | 2386 | **0.686** | 0.000 |
| A | 15 | **4.00%** | 4164 | 66716 | 2782 | **0.741** | 0.000 | 47816 | 1984 | **0.957** | 0.000 |
| A | 15 | **4.50%** | 4685 | 54131 | 2612 | **0.923** | 0.000 | 28954 | 1399 | **1.000** | 0.000 |
| A | 15 | **5.00%** | 5206 | 42338 | 2366 | **0.992** | 0.000 | 15110 | 850 | **1.000** | 0.002 |
| A | 15 | **5.50%** | 5726 | 30191 | 1962 | **0.999** | 0.000 | 6455 | 423 | **1.000** | **0.232** |
| A | 15 | **6.00%** | 6247 | 20094 | 1539 | **1.000** | 0.001 | 2133 | 166 | **1.000** | **0.995** |
| N-A | - | - | 500 | 111053 | 500 | 0.128 | 0.000 | 105609 | 500 | 0.12 | 0.000 |
| N-A | - | - | 1000 | 108364 | 1000 | 0.141 | 0.000 | 102169 | 1000 | 0.143 | 0.000 |
| N-A | - | - | 1500 | 105205 | 1500 | 0.184 | 0.000 | 98262 | 1500 | 0.197 | 0.000 |
| N-A | - | - | **2000** | 99160 | 2000 | **0.258** | 0.000 | 88296 | 2000 | **0.306** | 0.02 |
| N-A | - | - | **2500** | 90766 | 2500 | **0.353** | 0.000 | 69418 | 2500 | **0.478** | **0.142** |
| N-A | - | - | **3000** | 79890 | 3000 | **0.487** | 0.024 | 45115 | 3000 | **0.69** | **0.39** |
| N-A | - | - | **3500** | 63235 | 3500 | **0.644** | **0.084** | 19906 | 3500 | **0.872** | **0.697** |

Table S3. Summary of theta-logistic model output considering 44 harvest scenarios under the intermediate_245 climate/disturbance scenario. H_type (harvest type) is either A (adaptive scenarios) or N-A (non-adaptive scenarios); M_int is the monitoring interval; H is the percentage of the independent-aged female Pacific walrus population that is harvested annually in each scenario; A_t_ is the absolute harvest under each scenario; N_t_ is the mean estimated population size at each timestep of the simulation (i.e., the years 2050 and 2100); MO1 and MO2 are the probabilities of meeting Management Objective 1 (sustainability) and Management Objective 2 (avoiding quasi-extinction). Simulations which fail either management objective on at least one criterion according to the pre-defined risk tolerance thresholds (25% for MO1, 5% for MO2) are bolded.

| **H_type** | **M_int** | **H** | **A_2020_** | **N_2050_** | **A_2050_** | **MO1_2050_** | **MO2_2050_** | **N_2100_** | **A_2100_** | **MO1_2100_** | **MO2_2100_** |
| --- | --- | --- | --- | --- | --- | --- | --- | --- | --- | --- | --- |
| - | - | 0.00% | 0 | 84027 | 0 | 0.054 | 0.000 | 74234 | 0 | 0.083 | 0.000 |
| A | 5 | 0.50% | 520 | 82856 | 414 | 0.075 | 0.000 | 73065 | 365 | 0.083 | 0.000 |
| A | 5 | 1.00% | 1041 | 81316 | 813 | 0.078 | 0.000 | 70561 | 705 | 0.118 | 0.000 |
| A | 5 | 1.50% | 1561 | 78086 | 1171 | 0.107 | 0.000 | 66923 | 1003 | 0.172 | 0.000 |
| A | 5 | 2.00% | 2082 | 77069 | 1541 | 0.114 | 0.000 | 64619 | 1292 | 0.199 | 0.000 |
| A | 5 | **2.50%** | 2603 | 73687 | 1842 | 0.136 | 0.000 | 59242 | 1481 | **0.305** | 0.000 |
| A | 5 | **3.00%** | 3123 | 70890 | 2126 | 0.176 | 0.000 | 54051 | 1621 | **0.448** | 0.000 |
| A | 5 | **3.50%** | 3644 | 66644 | 2332 | 0.249 | 0.000 | 46932 | 1642 | **0.683** | 0.000 |
| A | 5 | **4.00%** | 4164 | 61059 | 2442 | **0.375** | 0.000 | 38086 | 1523 | **0.906** | 0.000 |
| A | 5 | **4.50%** | 4685 | 54889 | 2470 | **0.530** | 0.000 | 28557 | 1285 | **0.993** | 0.000 |
| A | 5 | **5.00%** | 5206 | 48439 | 2421 | **0.704** | 0.000 | 20161 | 1008 | **1.000** | 0.000 |
| A | 5 | **5.50%** | 5726 | 40810 | 2244 | **0.907** | 0.000 | 13107 | 720 | **1.000** | 0.000 |
| A | 5 | **6.00%** | 6247 | 34195 | 2051 | **0.977** | 0.000 | 8274 | 496 | **1.000** | 0.022 |
| A | 10 | 0.50% | 520 | 83843 | 445 | 0.063 | 0.000 | 73676 | 368 | 0.077 | 0.000 |
| A | 10 | 1.00% | 1041 | 81314 | 865 | 0.063 | 0.000 | 70178 | 701 | 0.108 | 0.000 |
| A | 10 | 1.50% | 1561 | 79697 | 1274 | 0.08 | 0.000 | 67948 | 1019 | 0.126 | 0.000 |
| A | 10 | 2.00% | 2082 | 76029 | 1618 | 0.120 | 0.000 | 63279 | 1265 | 0.213 | 0.000 |
| A | 10 | **2.50%** | 2603 | 73409 | 1957 | 0.142 | 0.000 | 59374 | 1484 | **0.315** | 0.000 |
| A | 10 | **3.00%** | 3123 | 69726 | 2227 | 0.197 | 0.000 | 53255 | 1597 | **0.479** | 0.000 |
| A | 10 | **3.50%** | 3644 | 65308 | 2441 | **0.294** | 0.000 | 45550 | 1594 | **0.740** | 0.000 |
| A | 10 | **4.00%** | 4164 | 58975 | 2538 | **0.431** | 0.000 | 35483 | 1419 | **0.945** | 0.000 |
| A | 10 | **4.50%** | 4685 | 52535 | 2580 | **0.596** | 0.000 | 25380 | 1142 | **1.000** | 0.000 |
| A | 10 | **5.00%** | 5206 | 43679 | 2438 | **0.836** | 0.000 | 15830 | 791 | **1.000** | 0.000 |
| A | 10 | **5.50%** | 5726 | 35402 | 2246 | **0.957** | 0.000 | 9035 | 496 | **1.000** | 0.021 |
| A | 10 | **6.00%** | 6247 | 27446 | 1978 | **0.999** | 0.000 | 4671 | 280 | **1.000** | **0.572** |
| **H_type** | **M_int** | **H** | **A_2020_** | **N_2050_** | **A_2050_** | **MO1_2050_** | **MO2_2050_** | **N_2100_** | **A_2100_** | **MO1_2100_** | **MO2_2100_** |
| A | 15 | 0.50% | 520 | 84025 | 447 | 0.066 | 0.000 | 73646 | 370 | 0.081 | 0.000 |
| A | 15 | 1.00% | 1041 | 80693 | 858 | 0.082 | 0.000 | 69920 | 704 | 0.113 | 0.000 |
| A | 15 | 1.50% | 1561 | 78368 | 1251 | 0.098 | 0.000 | 66948 | 1011 | 0.147 | 0.000 |
| A | 15 | 2.00% | 2082 | 75708 | 1607 | 0.121 | 0.000 | 63460 | 1280 | 0.216 | 0.000 |
| A | 15 | **2.50%** | 2603 | 71849 | 1912 | 0.155 | 0.000 | 58159 | 1470 | **0.335** | 0.000 |
| A | 15 | **3.00%** | 3123 | 68347 | 2178 | 0.233 | 0.000 | 52307 | 1597 | **0.521** | 0.000 |
| A | 15 | **3.50%** | 3644 | 63514 | 2371 | **0.33** | 0.000 | 43880 | 1584 | **0.778** | 0.000 |
| A | 15 | **4.00%** | 4164 | 56816 | 2449 | **0.473** | 0.000 | 32979 | 1393 | **0.967** | 0.000 |
| A | 15 | **4.50%** | 4685 | 47579 | 2355 | **0.746** | 0.000 | 20897 | 1030 | **1.000** | 0.000 |
| A | 15 | **5.00%** | 5206 | 37851 | 2156 | **0.939** | 0.000 | 11199 | 644 | **1.000** | 0.012 |
| A | 15 | **5.50%** | 5726 | 27852 | 1842 | **0.993** | 0.000 | 4829 | 325 | **1.000** | **0.55** |
| A | 15 | **6.00%** | 6247 | 18221 | 1424 | **1.000** | 0.000 | 1486 | 120 | **1.000** | **0.999** |
| N-A | - | - | 500 | 83582 | 500 | 0.071 | 0.000 | 72548 | 500 | 0.105 | 0.000 |
| N-A | - | - | 1000 | 80238 | 1000 | 0.100 | 0.000 | 67370 | 1000 | 0.185 | 0.000 |
| N-A | - | - | **1500** | 78652 | 1500 | 0.110 | 0.000 | 60126 | 1500 | **0.321** | 0.013 |
| N-A | - | - | **2000** | 75535 | 2000 | 0.163 | 0.000 | 43778 | 2000 | **0.576** | **0.134** |
| N-A | - | - | **2500** | 68886 | 2500 | **0.252** | 0.003 | 20957 | 2500 | **0.833** | **0.479** |
| N-A | - | - | **3000** | 58929 | 3000 | **0.422** | **0.023** | 6656 | 3000 | **0.956** | **0.799** |
| N-A | - | - | **3500** | 46470 | 3500 | **0.593** | **0.131** | 2018 | 3500 | **0.983** | **0.943** |

Table S4. Summary of theta-logistic model output considering 44 harvest scenarios under the intermediate_585 climate/disturbance scenario. H_type (harvest type) is either A (adaptive scenarios) or N-A (non-adaptive scenarios); M_int is the monitoring interval; H is the percentage of the independent-aged female Pacific walrus population that is harvested annually in each scenario; A_t_ is the absoluate harvest under each scenario; N_t_ is the mean estimated population size at each timestep of the simulation (i.e., the years 2050 and 2100); ; MO1 and MO2 are the probabilities of meeting Management Objective 1 (sustainability) and Management Objective 2 (avoiding quasi-extinction). Simulations which fail either management objective on at least one criterion according to the pre-defined risk tolerance thresholds (25% for MO1, 5% for MO2) are bolded.

| **H_type** | **M_int** | **H** | **A_2020_** | **N_2050_** | **A_2050_** | **MO1_2050_** | **MO2_2050_** | **N_2100_** | **A_2100_** | **MO1_2100_** | **MO2_2100_** |
| --- | --- | --- | --- | --- | --- | --- | --- | --- | --- | --- | --- |
| - | - | 0.00% | 0 | 83471 | 0 | 0.054 | 0.000 | 63685 | 0 | 0.06 | 0.000 |
| A | 5 | 0.50% | 520 | 81932 | 409 | 0.075 | 0.000 | 62219 | 311 | 0.077 | 0.000 |
| A | 5 | 1.00% | 1041 | 79849 | 798 | 0.089 | 0.000 | 60012 | 600 | 0.091 | 0.000 |
| A | 5 | 1.50% | 1561 | 78595 | 1178 | 0.102 | 0.000 | 58611 | 879 | 0.116 | 0.000 |
| A | 5 | 2.00% | 2082 | 75925 | 1518 | 0.098 | 0.000 | 55595 | 1111 | 0.151 | 0.000 |
| A | 5 | 2.50% | 2603 | 73587 | 1839 | 0.125 | 0.000 | 52296 | 1307 | 0.232 | 0.000 |
| A | 5 | **3.00%** | 3123 | 70307 | 2109 | 0.165 | 0.000 | 47861 | 1435 | **0.356** | 0.000 |
| A | 5 | **3.50%** | 3644 | 64946 | 2273 | **0.268** | 0.000 | 41314 | 1445 | **0.599** | 0.000 |
| A | 5 | **4.00%** | 4164 | 59834 | 2393 | **0.367** | 0.000 | 33851 | 1354 | **0.869** | 0.000 |
| A | 5 | **4.50%** | 4685 | 53352 | 2400 | **0.558** | 0.000 | 25457 | 1145 | **0.985** | 0.000 |
| A | 5 | **5.00%** | 5206 | 46264 | 2313 | **0.754** | 0.000 | 17613 | 880 | **1.000** | 0.000 |
| A | 5 | **5.50%** | 5726 | 39821 | 2190 | **0.908** | 0.000 | 11693 | 643 | **1.000** | 0.000 |
| A | 5 | **6.00%** | 6247 | 33529 | 2011 | **0.979** | 0.000 | 7401 | 444 | **1.000** | 0.000 |
| A | 10 | 0.50% | 520 | 82215 | 439 | 0.058 | 0.000 | 62368 | 311 | 0.064 | 0.000 |
| A | 10 | 1.00% | 1041 | 80330 | 858 | 0.063 | 0.000 | 60325 | 603 | 0.086 | 0.000 |
| A | 10 | 1.50% | 1561 | 78669 | 1262 | 0.089 | 0.000 | 58229 | 873 | 0.097 | 0.000 |
| A | 10 | 2.00% | 2082 | 75428 | 1612 | 0.114 | 0.000 | 55392 | 1107 | 0.163 | 0.000 |
| A | 10 | 2.50% | 2603 | 72808 | 1946 | 0.123 | 0.000 | 51452 | 1286 | 0.247 | 0.000 |
| A | 10 | **3.00%** | 3123 | 68651 | 2201 | 0.188 | 0.000 | 46585 | 1397 | **0.383** | 0.000 |
| A | 10 | **3.50%** | 3644 | 63730 | 2395 | **0.269** | 0.000 | 39881 | 1395 | **0.660** | 0.000 |
| A | 10 | **4.00%** | 4164 | 57306 | 2478 | **0.443** | 0.000 | 31270 | 1250 | **0.924** | 0.000 |
| A | 10 | **4.50%** | 4685 | 50550 | 2499 | **0.663** | 0.000 | 22152 | 996 | **0.993** | 0.000 |
| A | 10 | **5.00%** | 5206 | 42677 | 2401 | **0.851** | 0.000 | 13915 | 695 | **1.000** | 0.000 |
| A | 10 | **5.50%** | 5726 | 34195 | 2188 | **0.969** | 0.000 | 7800 | 429 | **1.000** | 0.045 |
| A | 10 | **6.00%** | 6247 | 26236 | 1909 | **0.998** | 0.000 | 3959 | 237 | **1.000** | **0.77** |
| **H_type** | **M_int** | **H** | **A_2020_** | **N_2050_** | **A_2050_** | **MO1_2050_** | **MO2_2050_** | **N_2100_** | **A_2100_** | **MO1_2100_** | **MO2_2100_** |
| A | 15 | 0.50% | 520 | 82047 | 438 | 0.073 | 0.000 | 62292 | 319 | 0.069 | 0.000 |
| A | 15 | 1.00% | 1041 | 79866 | 853 | 0.061 | 0.000 | 60479 | 620 | 0.099 | 0.000 |
| A | 15 | 1.50% | 1561 | 77285 | 1237 | 0.092 | 0.000 | 57477 | 885 | 0.136 | 0.000 |
| A | 15 | 2.00% | 2082 | 75381 | 1607 | 0.113 | 0.000 | 55053 | 1132 | 0.166 | 0.000 |
| A | 15 | **2.50%** | 2603 | 72025 | 1921 | 0.144 | 0.000 | 51103 | 1316 | **0.260** | 0.000 |
| A | 15 | **3.00%** | 3123 | 67666 | 2167 | 0.218 | 0.000 | 46001 | 1428 | **0.413** | 0.000 |
| A | 15 | **3.50%** | 3644 | 61805 | 2319 | 0.330 | 0.000 | 38063 | 1395 | **0.732** | 0.000 |
| A | 15 | **4.00%** | 4164 | 54729 | 2372 | 0.525 | 0.000 | 28493 | 1221 | **0.964** | 0.000 |
| A | 15 | **4.50%** | 4685 | 46057 | 2298 | 0.778 | 0.000 | 17995 | 900 | **0.999** | 0.000 |
| A | 15 | **5.00%** | 5206 | 36606 | 2106 | 0.930 | 0.000 | 9479 | 554 | **1.000** | 0.032 |
| A | 15 | **5.50%** | 5726 | 26347 | 1763 | 0.996 | 0.000 | 3907 | 269 | **1.000** | **0.785** |
| A | 15 | **6.00%** | 6247 | 17272 | 1369 | 1.000 | 0.000 | 1145 | 95 | **1.000** | **1.000** |
| N-A | - | - | 500 | 81253 | 500 | 0.065 | 0.000 | 60480 | 500 | 0.101 | 0.000 |
| N-A | - | - | 1000 | 79099 | 1000 | 0.091 | 0.000 | 56823 | 1000 | 0.176 | 0.000 |
| N-A | - | - | **1500** | 76613 | 1500 | 0.121 | 0.000 | 48483 | 1500 | **0.373** | 0.018 |
| N-A | - | - | **2000** | 74017 | 2000 | 0.161 | 0.002 | 33397 | 2000 | **0.630** | **0.184** |
| N-A | - | - | **2500** | 66895 | 2500 | **0.290** | 0.004 | 14686 | 2500 | **0.859** | **0.584** |
| N-A | - | - | **3000** | 56838 | 3000 | **0.433** | 0.028 | 4233 | 3000 | **0.960** | **0.852** |
| N-A | - | - | **3500** | 44455 | 3500 | **0.601** | **0.11** | 714 | 3500 | **0.995** | **0.974** |

Table S5. Summary of theta-logistic model output considering 44 harvest scenarios under the pessimistic climate/disturbance scenario. H_type (harvest type) is either A (adaptive scenarios) or N-A (non-adaptive scenarios); M_int is the monitoring interval; H is the percentage of the independent-aged female Pacific walrus population that is harvested annually in each scenario; A_t_ is the absolute harvest under each scenario; N_t_ is the mean estimated population size at each timestep of the simulation (i.e., the years 2050 and 2100); MO1 and MO2 are the probabilities of meeting Management Objective 1 (sustainability) and Management Objective 2 (avoiding quasi-extinction). Simulations which fail either management objective on at least one criterion according to the pre-defined risk tolerance thresholds (25% for MO1, 5% for MO2) are bolded.

| **H_type** | **M_int** | **H** | **A_2020_** | **N_2050_** | **A_2050_** | **MO1_2050_** | **MO2_2050_** | **N_2100_** | **A_2100_** | **MO1_2100_** | **MO2_2100_** |
| --- | --- | --- | --- | --- | --- | --- | --- | --- | --- | --- | --- |
| - | - | 0.00% | 0 | 59422 | 0 | 0.031 | 0.000 | 42812 | 0 | 0.067 | 0.000 |
| A | 5 | 0.50% | 520 | 58311 | 291 | 0.036 | 0.000 | 41304 | 206 | 0.084 | 0.000 |
| A | 5 | 1.00% | 1041 | 57752 | 577 | 0.036 | 0.000 | 40097 | 400 | 0.096 | 0.000 |
| A | 5 | 1.50% | 1561 | 57037 | 855 | 0.033 | 0.000 | 38194 | 572 | 0.143 | 0.000 |
| A | 5 | 2.00% | 2082 | 55457 | 1109 | 0.047 | 0.000 | 35748 | 714 | 0.206 | 0.000 |
| A | 5 | **2.50%** | 2603 | 54260 | 1356 | 0.063 | 0.000 | 32858 | 821 | **0.317** | 0.000 |
| A | 5 | **3.00%** | 3123 | 52813 | 1584 | 0.068 | 0.000 | 29553 | 886 | **0.51** | 0.000 |
| A | 5 | **3.50%** | 3644 | 50989 | 1784 | 0.082 | 0.000 | 25561 | 894 | **0.725** | 0.000 |
| A | 5 | **4.00%** | 4164 | 48060 | 1922 | 0.125 | 0.000 | 21072 | 842 | **0.929** | 0.000 |
| A | 5 | **4.50%** | 4685 | 44333 | 1994 | 0.203 | 0.000 | 16418 | 738 | **0.99** | 0.000 |
| A | 5 | **5.00%** | 5206 | 40115 | 2005 | **0.335** | 0.000 | 12071 | 603 | **1.000** | 0.000 |
| A | 5 | **5.50%** | 5726 | 35705 | 1963 | **0.541** | 0.000 | 8487 | 466 | **1.000** | 0.030 |
| A | 5 | **6.00%** | 6247 | 30393 | 1823 | **0.756** | 0.000 | 5533 | 331 | **1.000** | **0.331** |
| A | 10 | 0.50% | 520 | 58357 | 341 | 0.021 | 0.000 | 41429 | 207 | 0.072 | 0.000 |
| A | 10 | 1.00% | 1041 | 58111 | 679 | 0.022 | 0.000 | 40160 | 401 | 0.084 | 0.000 |
| A | 10 | 1.50% | 1561 | 56710 | 989 | 0.045 | 0.000 | 38111 | 571 | 0.139 | 0.000 |
| A | 10 | 2.00% | 2082 | 55256 | 1284 | 0.044 | 0.000 | 35524 | 710 | 0.199 | 0.000 |
| A | 10 | **2.50%** | 2603 | 54344 | 1578 | 0.046 | 0.000 | 32867 | 821 | **0.325** | 0.000 |
| A | 10 | **3.00%** | 3123 | 51877 | 1802 | 0.084 | 0.000 | 28600 | 858 | **0.543** | 0.000 |
| A | 10 | **3.50%** | 3644 | 50280 | 2033 | 0.094 | 0.000 | 24416 | 854 | **0.772** | 0.000 |
| A | 10 | **4.00%** | 4164 | 46573 | 2149 | 0.154 | 0.000 | 19147 | 765 | **0.959** | 0.000 |
| A | 10 | **4.50%** | 4685 | 41838 | 2175 | **0.307** | 0.000 | 13840 | 622 | **0.998** | 0.000 |
| A | 10 | **5.00%** | 5206 | 37307 | 2181 | **0.469** | 0.000 | 9368 | 468 | **1.000** | 0.012 |
| A | 10 | **5.50%** | 5726 | 30754 | 2023 | **0.757** | 0.000 | 5510 | 303 | **1.000** | **0.346** |
| A | 10 | **6.00%** | 6247 | 24414 | 1816 | **0.942** | 0.000 | 2902 | 174 | **1.000** | **0.961** |
| **H_type** | **M_int** | **H** | **A_2020_** | **N_2050_** | **A_2050_** | **MO1_2050_** | **MO2_2050_** | **N_2100_** | **A_2100_** | **MO1_2100_** | **MO2_2100_** |
| A | 15 | 0.50% | 520 | 57945 | 339 | 0.032 | 0.000 | 41053 | 209 | 0.080 | 0.000 |
| A | 15 | 1.00% | 1041 | 57884 | 675 | 0.020 | 0.000 | 39958 | 408 | 0.089 | 0.000 |
| A | 15 | 1.50% | 1561 | 56900 | 993 | 0.032 | 0.000 | 38302 | 586 | 0.136 | 0.000 |
| A | 15 | 2.00% | 2082 | 55261 | 1286 | 0.048 | 0.000 | 35464 | 726 | 0.204 | 0.000 |
| A | 15 | **2.50%** | 2603 | 53362 | 1542 | 0.061 | 0.000 | 31904 | 822 | **0.375** | 0.000 |
| A | 15 | **3.00%** | 3123 | 51446 | 1783 | 0.080 | 0.000 | 27729 | 867 | **0.609** | 0.000 |
| A | 15 | **3.50%** | 3644 | 48302 | 1949 | 0.145 | 0.000 | 22392 | 834 | **0.861** | 0.000 |
| A | 15 | **4.00%** | 4164 | 45176 | 2081 | 0.213 | 0.000 | 16934 | 744 | **0.991** | 0.000 |
| A | 15 | **4.50%** | 4685 | 39162 | 2044 | **0.417** | 0.000 | 11058 | 570 | **1.000** | 0.004 |
| A | 15 | **5.00%** | 5206 | 32132 | 1914 | **0.698** | 0.000 | 6060 | 367 | **1.000** | **0.282** |
| A | 15 | **5.50%** | 5726 | 24065 | 1655 | **0.932** | 0.001 | 2575 | 185 | **1.000** | **0.967** |
| A | 15 | **6.00%** | 6247 | 16179 | 1317 | **0.999** | 0.012 | 705 | 62 | **1.000** | **1.000** |
| N-A | - | - | 500 | 58001 | 500 | 0.040 | 0.000 | 38941 | 500 | 0.150 | 0.000 |
| N-A | - | - | **1000** | 56144 | 1000 | 0.042 | 0.000 | 29410 | 1000 | **0.476** | 0.033 |
| N-A | - | - | 1500 | 54299 | 1500 | 0.069 | 0.000 | 12113 | 1500 | **0.855** | **0.426** |
| N-A | - | - | 2000 | 51832 | 2000 | 0.133 | 0.003 | 2391 | 2000 | **0.973** | **0.861** |
| N-A | - | - | 2500 | 47739 | 2500 | 0.212 | 0.013 | 189 | 2500 | **0.997** | **0.989** |
| N-A | - | - | 3000 | 40031 | 3000 | **0.356** | **0.063** | 20 | 3000 | **1.000** | **0.996** |
| N-A | - | - | 3500 | 31144 | 3500 | **0.523** | **0.16** | 0 | 3500 | **1.000** | **1.000** |

Table S6. Sensitivity of the model to risk tolerance levels, displaying model output for the maximum percentage-based harvest allowable to meet MO1 with risk tolerance levels of 15% (low), 25% (intermediate), and 35% (high) for each of the four combined climate/disturbance scenarios to the end of the 21^st^ century. H_type (harvest type) is either A (adaptive scenarios) or N-A (non-adaptive scenarios); M_int is the monitoring interval; H is the percentage of the independent-aged female Pacific walrus population that is harvested annually in each scenario; A_t_ is the absolute harvest under each scenario; N_t_ is the mean estimated population size at each timestep of the simulation (i.e., the years 2050 and 2100); MO1 and MO2 are the probabilities of meeting Management Objective 1 (sustainability) and Management Objective 2 (avoiding quasi-extinction).

| Model Parameters | | | | | | Output – Timestep 30 (2050) | | | | Output – Timestep 80 (2100) | | | |
| --- | --- | --- | --- | --- | --- | --- | --- | --- | --- | --- | --- | --- | --- |
| Scenario | Risk Tolerance | H_type | M_int | H | A_2020_ | N_2050_ | A_2050_ | MO1_2050_ | MO2_2050_ | N_2100_ | A_2100_ | MO1_2100_ | MO2_2100_ |
| **Optimistic Scenario** | | | | | | | | | | | | | |
| OPT | 15% | A | 10 | 1.26% | 1311 | 106,406 | 1340 | 0.149 | 0.000 | 99,747 | 1253 | 0.149 | 0.000 |
| OPT | 25% | A | 10 | 2.27% | 2363 | 97,771 | 2219 | 0.223 | 0.000 | 90,399 | 2052 | 0.249 | 0.000 |
| OPT | 35% | A | 10 | 2.75% | 2863 | 92,540 | 2544 | 0.296 | 0.000 | 84,205 | 2315 | 0.349 | 0.000 |
| **Intermediate_245 Scenario** | | | | | | | | | | | | | |
| I_245 | 15% | A | 10 | 1.35% | 1405 | 85,234 | 1150 | 0.082 | 0.000 | 68,720 | 927 | 0.150 | 0.000 |
| I_245 | 25% | A | 10 | 2.3% | 2394 | 80,712 | 1856 | 0.117 | 0.000 | 61,930 | 1424 | 0.249 | 0.000 |
| I_245 | 35% | A | 10 | 2.70% | 2811 | 78,138 | 2199 | 0.150 | 0.000 | 58,158 | 1570 | 0.349 | 0.000 |
| **Intermediate_585 Scenario** | | | | | | | | | | | | | |
| I_585 | 15% | A | 10 | 1.80% | 1874 | 81,398 | 1465 | 0.100 | 0.000 | 55,758 | 1003 | 0.150 | 0.000 |
| I_585 | 25% | A | 10 | 2.50% | 2603 | 77,022 | 1925 | 0.131 | 0.000 | 51,013 | 1275 | 0.250 | 0.000 |
| I_585 | 35% | A | 10 | 2.90% | 3019 | 74,604 | 2163 | 0.185 | 0.000 | 47,879 | 1388 | 0.350 | 0.000 |
| **Pessimistic Scenario** | | | | | | | | | | | | | |
| PESS | 15% | A | 10 | 1.70% | 1770 | 66,360 | 1128 | 0.040 | 0.000 | 37,683 | 640 | 0.150 | 0.000 |
| PESS | 25% | A | 10 | 2.25% | 2342 | 63,413 | 1426 | 0.057 | 0.000 | 34,203 | 769 | 0.250 | 0.000 |
| PESS | 35% | A | 10 | 2.60% | 2707 | 62,776 | 1632 | 0.072 | 0.000 | 32,081 | 834 | 0.349 | 0.000 |

**Supplemental Figures:**

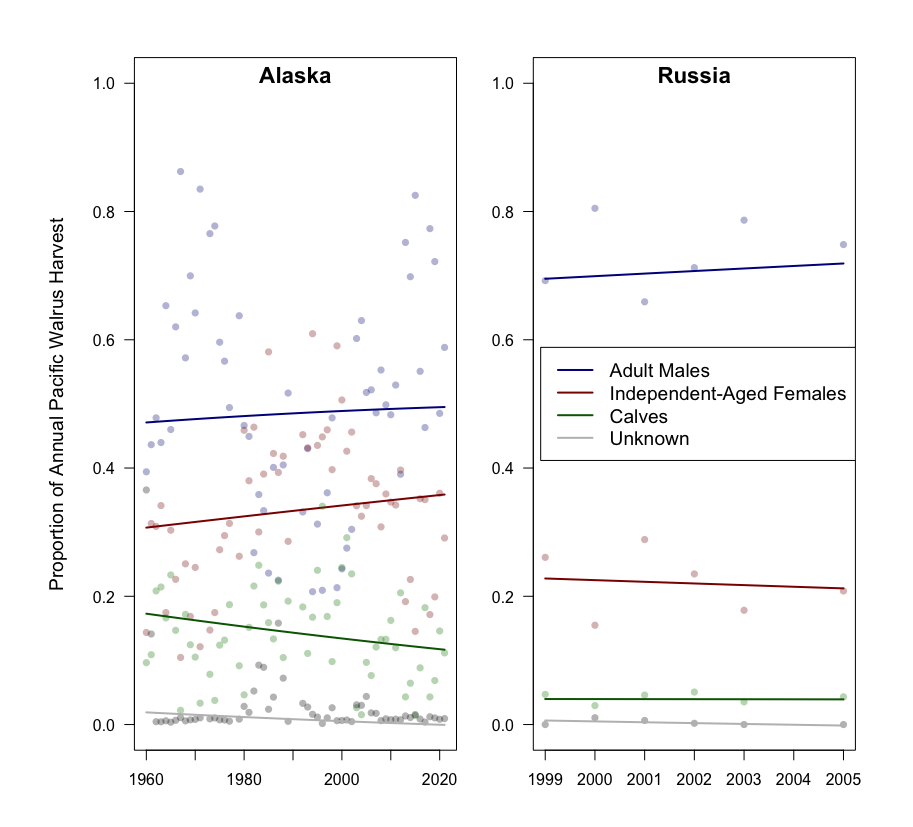
 Figure S1. Demographic harvest trends for Alaska and Russia. Lines indicate predicted output of Dirichlet regression models. No trends were significant in the Dirichlet regression analysis. Alaskan harvest trends are from FWS harvest monitoring programs (Garlich-Miller et al. 2006; USFWS 2024), and Russian trends are limited to a short timeframe (1999-2005) over which demographic harvest data were collected (Garlich-Miller et al. 2006).

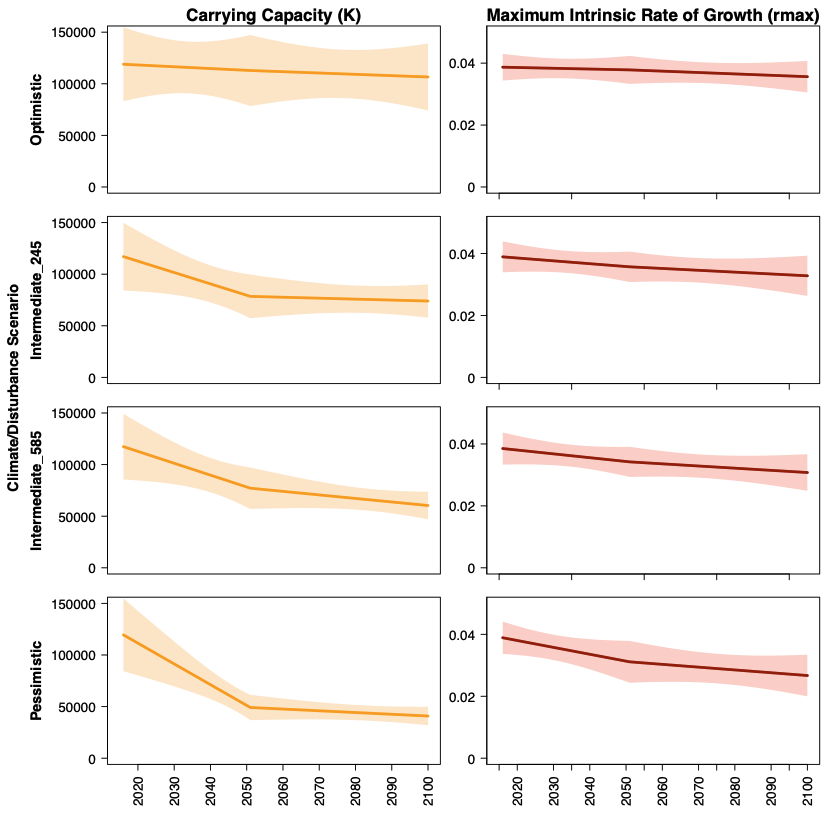

Figure S2. Output from the PCoD model (Johnson et al., 2023), indicating projected changes to *K_t_* and *r_max,t_* under the four climate-disturbance scenarios considered in this study (i.e., Table 1) to the end of the 21^st^ century. The optimistic scenario incorporates low levels of disturbance under an intermediate (ssp245) climate scenario, the intermediate_245 scenario incorporates intermediate levels of disturbance under an intermediate (ssp245) climate scenario, the intermediate_585 scenario incorporates intermediate levels of disturbance under a pessimistic (ssp585) climate scenario, and the pessimistic scenario incorporates high levels of disturbance under a pessimistic (ssp585) climate scenario. Shaded polygons show 95% confidence intervals from PCoD model output.

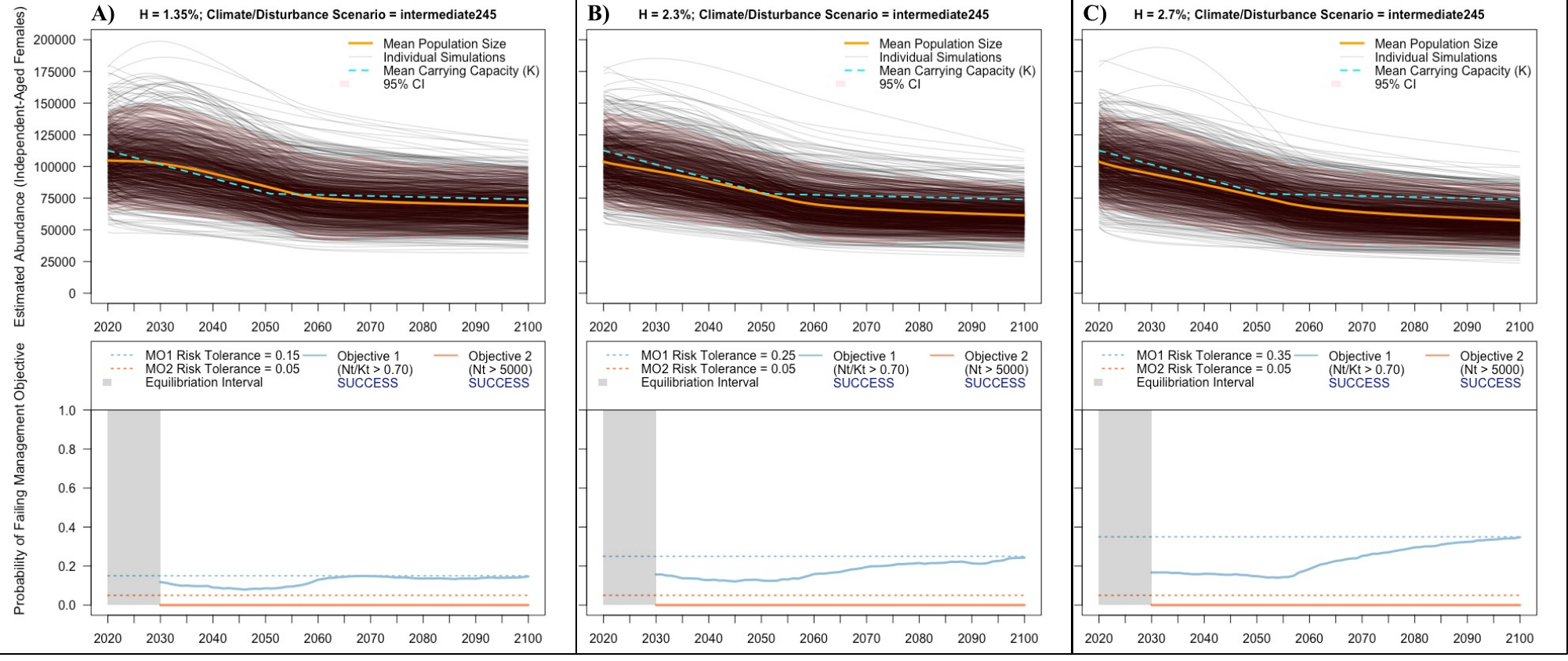

Figure S3. Example output from the theta-logistic model to visualize the sensitivity of MO1 (sustainability) to different risk tolerance levels, considering three adaptive harvest scenarios under the intermediate_245 combined climate/disturbance scenario and a 10-year (intermediate) monitoring interval projecting forward to the end of the 21^st^ century. Column A considers an annual harvest rate of 1.1% and a MO1 risk tolerance level of 15% (low), column B considers an annual harvest rate of 1.90% and a MO1 risk tolerance level of 25% (medium), and column C considers an annual harvest rate of 2.35% and a MO1 risk tolerance level of 35% (high). In the top panels, orange lines represent the mean estimated population size of independent-aged female Pacific walruses, gray lines are individual simulations (n=1000), cyan dashed lines represent mean carrying capacity estimates from each climate/disturbance scenario, and the red shaded region is a 95% credible interval of theta-logistic simulations. Bottom panels represent the associated probability of failing to meet risk tolerance thresholds for pre-defined management objectives (MO1 & MO2) at each timestep of the simulation.

**Supplemental Material – Works Cited**

Estes, J. A. and V. N. Gol'tsev. 1984. Abundance and distribution of the Pacific walrus Odobenus rosmarus divergens: results of the first Soviet-American joint aerial survey, autumn 1975. U.S. Department of Commerce, National Oceanic and Atmospheric Adminstration, National Marine Fisheries Service, Washington D.C.

Fedoseev, G. A. 1984. Present status of the population of walruses Odobenus rosmarus in the eastern Arctic and Bering Sea. Pages 73-85 in V.E. Rodin, A.S. Perlov, A.A. Berzin, G.M. Gavrilov, A.I. Shevchenko, N.S. Fadeev and E.B. Kucheriavenko eds. Marine mammals of the far east. TINRO, Vladivostok, Russia.

Fedoseev, G. A. and E. V. Razlivalov. 1986. Distribution and abundance of walruses in the eastern Arctic and Bering Sea in the Autumn of 1985. All-Union Research Institute of Marine Fisheries and Oceanography (VNIRO), Moscow, Russia.

Gilbert, J., G. Fedoseev, D. Seagars, E. Razlivalov and A. Lachugin. 1992. Aerial census of Pacific walrus 1990. U.S. Department of the Interior, U.S. Fish and Wildlife Service, Marine Mammals Management Administrative Report, MMM 92-1, Anchorage, AK.

Gilbert, J. R. 1989. Aerial census of Pacific walruses in the Chukchi Sea, 1985. Marine Mammal Science 5:17-28.

Johnson, A., J. Burns, W. Dusenberry and R. Jones. 1982. Aerial survey of Pacific walrus, 1980. U.S. Department of the Interior, U.S. Fish and Wildlife Service, Marine Mammals Management, Anchorage, Alaska.

National Marine Fisheries Service (NMFS). 2016. Guidelines for preparing stock assessment reports pursuant to the 1994 amendments to the Marine Mammal Protection Act. 23 p. Available online: https://www.fisheries.noaa.gov/national/marine-mammal-protection/guidelines-assessing-marine-mammal-stocks . Accessed June 2022.

Taylor, R. L. and M. S. Udevitz. 2015. Demography of the Pacific walrus (Odobenus rosmarus divergens): 1974–2006. Marine Mammal Science 31:231-254.

Udevitz, M. S., J. R. Gilbert and G. A. Fedoseev. 2001. Comparison of methods used to estimate numbers of walruses on sea ice. Marine Mammal Science 17:601–616.

USFWS (U.S. Fish and Wildlife Service). 2024. Pacific Walrus Marking, Tagging, and Reporting Program. USFWS Marine Mammals Management, Anchorage, AK, USA. Unpublished database.
